# A combination of cross-neutralizing antibodies synergizes to prevent SARS-CoV-2 and SARS-CoV pseudovirus infection

**DOI:** 10.1101/2021.02.11.430866

**Authors:** Hejun Liu, Meng Yuan, Deli Huang, Sandhya Bangaru, Chang-Chun D. Lee, Linghang Peng, Xueyong Zhu, David Nemazee, Marit J. van Gils, Rogier W. Sanders, Hans-Christian Kornau, S. Momsen Reincke, Harald Prüss, Jakob Kreye, Nicholas C. Wu, Andrew B. Ward, Ian A. Wilson

**Affiliations:** Department of Integrative Structural and Computational Biology, The Scripps Research Institute, La Jolla, CA 92037, USA; Department of Immunology and Microbiology and Infection Prevention, The Scripps Research Institute, La Jolla, CA 92037, USA; Department of Medical Microbiology, Amsterdam University Medical Centers, Location AMC, University of Amsterdam, Amsterdam, The Netherlands; Department of Microbiology and Immunology, Weill Medical College of Cornell University, New York, NY 10021, USA; German Center for Neurodegenerative Diseases (DZNE) Berlin, Berlin, Germany; Neuroscience Research Center (NWFZ), Cluster NeuroCure, Charité-Universitätsmedizin Berlin, corporate member of Freie Universität Berlin, Humboldt-Universität Berlin, and Berlin Institute of Health, Berlin, Germany; Helmholtz Innovation Lab BaoBab, Berlin, Germany; Department of Neurology and Experimental Neurology, Charité-Universitätsmedizin Berlin, corporate member of Freie Universität Berlin, Humboldt-Universität Berlin, and Berlin Institute of Health, Berlin, Germany; Department of Pediatric Neurology, Charité-Universitätsmedizin Berlin, corporate member of Freie Universität Berlin, Humboldt-Universität Berlin, and Berlin Institute of Health, Berlin, Germany; Department of Biochemistry, University of Illinois at Urbana-Champaign, Urbana, IL 61801, USA; Carl R. Woese Institute for Genomic Biology, University of Illinois at Urbana-Champaign, Urbana, IL 61801, USA; Center for Biophysics and Quantitative Biology, University of Illinois at Urbana-Champaign, Urbana, IL 61801, USA; The Skaggs Institute for Chemical Biology, The Scripps Research Institute, La Jolla, CA, 92037, USA

## Abstract

Coronaviruses have caused several epidemics and pandemics including the ongoing coronavirus disease 2019 (COVID-19). Some prophylactic vaccines and therapeutic antibodies have already showed striking effectiveness against COVID-19. Nevertheless, concerns remain about antigenic drift in SARS-CoV-2 as well as threats from other sarbecoviruses. Cross-neutralizing antibodies to SARS-related viruses provide opportunities to address such concerns. Here, we report on crystal structures of a cross-neutralizing antibody CV38-142 in complex with the receptor binding domains from SARS-CoV-2 and SARS-CoV. Our structural findings provide mechanistic insights into how this antibody can accommodate antigenic variation in these viruses. CV38-142 synergizes with other cross-neutralizing antibodies, in particular COVA1-16, to enhance neutralization of SARS-CoV-2 and SARS-CoV. Overall, this study provides valuable information for vaccine and therapeutic design to address current and future antigenic drift in SARS-CoV-2 and to protect against zoonotic coronaviruses.

## INTRODUCTION

Severe acute respiratory syndrome coronavirus (SARS-CoV), middle east respiratory syndrome coronavirus (MERS-CoV) and SARS-CoV-2, have caused epidemics in the past two decades including the current pandemic of coronavirus disease 2019 (COVID-19). SARS-CoV-2 has already resulted in more than 100 million reported cases and almost 2.3 million deaths worldwide as of the beginning of February 2021 (https://covid19.who.int). Although these viruses have devastating consequences in the human population, they are of animal origin and have less morbidity or even no symptoms in their animal hosts (Cui et al., 2019; Tortorici and Veesler, 2019; Ye et al., 2020). In addition to these human β-coronaviruses (SARS-CoV, MERS-CoV, and SARS-CoV-2), other SARS-related coronaviruses (SARSr-CoVs) of the sarbecovirus subgenus within the β-coronavirus genus are found in mammalian reservoirs, such as bats and pangolins, and could also constitute potential pandemic threats to human health (Hu et al., 2015; Lam et al., 2020; Wacharapluesadee et al., 2021; Ye et al., 2020). Recently, mutations in SARS-CoV-2 were identified in farmed mink and these viruses were found to be reciprocally transmissible between humans and farmed mink (Welkers et al., 2021), further underscoring concerns about the long-term efficacy of current antibody therapies and vaccines under development (Mallapaty, 2020). Hence, identification and characterization of cross-neutralizing antibodies within the sarbecovirus subgenus are of value for design and development of therapeutics and next generation vaccines to mitigate against antigenic drift as well as future SARSr-CoV transmission to humans from the mammalian reservoir.

Since the spike protein is the major surface protein on sarbecoviruses, neutralizing antibodies are targeted towards the spike and many of these antibodies are able to prevent virus interaction with the host receptor, angiotensin-converting enzyme 2 (ACE2) (Piccoli et al., 2020; Yuan et al., 2020b). Other inhibition mechanisms also seem to be possible and are being assessed for other subsets of antibodies (Hansen et al., 2020; Piccoli et al., 2020; Pinto et al., 2020). The receptor binding domain (RBD) of the spike protein is highly immunogenic and can induce highly specific and potent neutralizing antibodies (nAbs) against SARS-CoV-2 virus (Barnes et al., 2020a; Barnes et al., 2020b; Brouwer et al., 2020; Cao et al., 2020; Ju et al., 2020; Kreye et al., 2020; Piccoli et al., 2020; Robbiani et al., 2020; Rogers et al., 2020; Yuan et al., 2020a; Zost et al., 2020). Many of these nAbs bind to the receptor binding site (RBS) on the RBD (Yuan et al., 2020b). However, the breadth of these nAbs is limited as the RBS shares relatively low sequence identity among sarbecoviruses; the RBS is only 48% conserved between SARS-CoV-2 and SARS-CoV compared to 73% for the complete RBD (84% identity for non-RBS regions of the RBD). The RBS is also prone to naturally occurring mutations, similar to the N-terminal domain (NTD), where insertions and deletions have also been found (Greaney et al., 2021b; Kemp et al., 2021; Liu et al., 2021; McCarthy et al., 2020; Starr et al., 2020; Tegally et al., 2020; Van Egeren et al., 2020; Voloch et al., 2020). Recent studies showed that many potent monoclonal neutralizing antibodies are subject to the antigenic drift or mutation on the RBD of the spike protein (Thomson et al., 2020; Wang et al., 2021a; Wang et al., 2021b; Weisblum et al., 2020; Wibmer et al., 2021), as well as polyclonal sera from convalescent or vaccinated individuals (Andreano et al., 2020; Greaney et al., 2021a; Liu et al., 2021; Wang et al., 2021a; Wang et al., 2021b; Weisblum et al., 2020; Wu et al., 2021).

We and others have reported cross-neutralizing antibodies that bind to a highly conserved cryptic site in receptor binding domain (RBD) of the spike (Liu et al., 2020; Lv et al., 2020; Yuan et al., 2020b; Zhou et al., 2020). Although the epitopes of these antibodies do not overlap with the ACE2 receptor binding site, some can sterically block ACE2 binding to the RBD or attenuate ACE2 binding affinity (Liu et al., 2020; Lv et al., 2020). Other RBD surfaces are also possible targets for cross-neutralizing antibodies, but are only moderately conserved within sarbecoviruses, although more so than the RBS. Such a site was originally identified as the epitope for antibody S309, which was isolated from a SARS patient, but cross-neutralizes SARS-CoV-2. S309 binds to a non-RBS surface containing an N-glycosylation site at N343 (Pinto et al., 2020). Further investigation is ongoing as to whether the S309 site is a common target for antibodies elicited by SARS-CoV-2 infection. Here, we report on cross-neutralization of sarbecoviruses by an *IGHV5-51* encoded antibody isolated from a SARS-CoV-2 patient. High-resolution crystal structures of CV38-142 were determined in complex with both SARS-CoV RBD and SARS-CoV-2 RBD in combination with another cross-neutralizing antibody COVA1-16. The structural information, along with binding data, revealed that CV38-142 can be combined with cross-neutralizing antibodies to other epitopes to generate therapeutic cocktails that to protect against SARS-CoV-2 variants, escape mutants, and future zoonotic coronavirus epidemics. The information may also inform next generation vaccine and therapeutic design (Barnes et al., 2020a).

## RESULTS

### CV38-142 neutralizes both SARS-CoV-2 and SARS-CoV pseudoviruses

Previously, we reported that antibody CV38-142 isolated from a COVID-19 patient showed potent neutralization on authentic SARS-CoV-2 virus (Munich isolate 984) and was able to cross-react with SARS-CoV (Kreye et al., 2020). CV38-142 is an *IGHV5-51-encoded* antibody with little somatic hypermutation (only four mutations in the amino-acid sequence). This germline heavy-chain gene was also used in another cross-reactive antibody CR3022 (Yuan et al., 2020c) that was isolated from a SARS patient (ter Meulen et al., 2006), but their CDR H3s are quite distinct. A biolayer interferometry (BLI) binding assay revealed that CV38-142 binds with high affinity not only to SARS-CoV-2 RBD (29 nM), but also SARS-CoV, RaTG13 and Guangdong pangolin coronavirus RBDs with roughly comparable affinity (36-99 nM) (Figure 1A). A pseudovirus neutralization assay showed that CV38-142 IgG neutralizes both SARS-CoV-2 and SARS-CoV with similar potency (3.5 and 1.4 μg/ml) (Figure 1B). Of note, the CV38-142 Fab exhibits much weaker or no neutralization in the same assay, which suggests that the avidity of bivalent CV38-142 IgG plays a crucial role in the neutralization (Figure 1B) as we also observed in other antibodies such as COVA1-16 (Liu et al., 2020).

**Figure 1.**
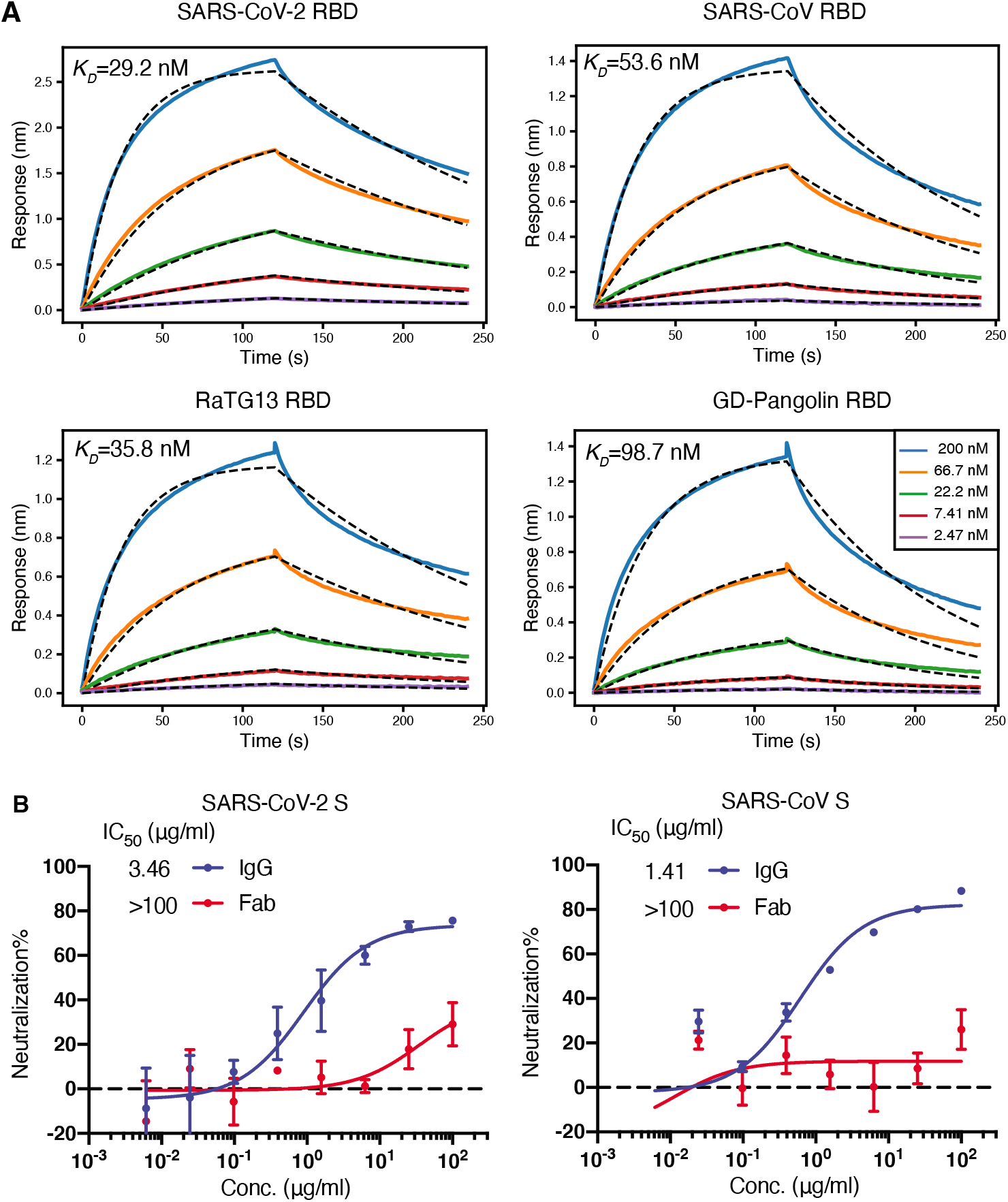
CV38-142 binds and cross-neutralizes SARS-CoV-2 and SARS-CoV. **A.** CV38-142 Fab binds to RBDs from human, bat and pangolin sarbecoviruses with generally similar affinities. Binding kinetics were measured by biolayer interferometry (BLI) with RBDs on the biosensor and Fab in solution. Concentrations of Fab serial dilution are shown in the middle insert panel. The association and disassociation were recorded in real time (s) on the x axis with binding response (nm) on the y axis with colored lines. Disassociation constant (K_D_) values were obtained by fitting a 1:1 binding model. The fitted curves are represented by the dash lines (black). **B.** CV38-142 neutralizes both SARS-CoV-2 and SARS-CoV, while its Fab counterpart barely neutralizes the two pseudotype viruses at the highest concentrations tested in the same neutralization assay. The IgG half-maximal inhibitory concentration (IC_50_) values (3.46 μg/ml for SARS-CoV-2 and 1.41 μg/mL for SARS-CoV) were determined using Prism software (version 8.4.3). Error bars indicate standard deviation (SD) of at least two biological replicates.

### CV38-142 can be combined with either RBS or CR3022 cryptic site antibodies

Recent reports on SARS-CoV-2 mutations in both human and mink populations give rise to concerns about viral escape from current vaccines and therapeutics in development (Andreano et al., 2020; Greaney et al., 2021a; Kemp et al., 2021; Mallapaty, 2020; Oude Munnink et al., 2021; Tegally et al., 2020; Voloch et al., 2020). However, antibody cocktails that bind to distinct epitopes can increase neutralization breadth and may help prevent escape mutations (Baum et al., 2020; Du et al., 2020; Greaney et al., 2021b; Hansen et al., 2020; Koenig et al., 2021). We previously reported that CV38-142 does not compete for RBD binding with other potent antibodies in our sample set, which are encoded by diverse germline genes, such as CV07-200 (IGHV1-2), CV07-209 (IGHV3-11), CV07-222 (IGHV1-2), CV07-250 (IGHV1-18), CV07-262 (IGHV1-2), CV38-113 (IGHV3-53), and CV38-183 (IGHV3-53) (Kreye et al., 2020). Here, we show that CV38-142 can bind either SARS-CoV-2 RBD or spike protein at the same time in a sandwich assay as CC12.1 and COVA2-39 (Figure 2A), which are potent IGHV3-53 neutralizing antibodies from different cohorts (Brouwer et al., 2020; Rogers et al., 2020). Since CC12.1 (Yuan et al., 2020a), as well as COVA2-39 (Wu et al., 2020) and CV07-250 (Kreye et al., 2020), bind to the RBS, these data suggest that CV38-142 can be combined with potent RBS antibodies derived from diverse germlines in an antibody cocktail. Hence, we tested whether CV38-142 could bind RBD at the same time as two other potent cross-neutralizing antibodies that target other sites on the RBD (Yuan et al., 2020b). The sandwich binding assay revealed that CV38-142 competes with S309 from a SARS patient (Pinto et al., 2020), but is compatible with COVA1-16, a cross neutralizing antibody to the CR3022 site isolated from a COVID-19 patient (Figure 2A) (Brouwer et al., 2020). We then assembled a cocktail consisting of different amounts and ratios of CV38-142 and COVA1-16. The cocktail showed enhanced potency in the 2D neutralization matrix assay with SARS-CoV-2 and enhanced potency and improved efficacy with SARS-CoV pseudoviruses, demonstrating that CV38-142 is a promising candidate for pairing with cross-neutralizing antibodies to the CR3022 cryptic site (Figure 2B). For example, 100% inhibition in the neutralization assay could be achieved with 1.6 μg/ml of each of CV138-142 and COVA1-16 with SARS-CoV-2 compared to >200 μg and 40 μg/ml for the individual antibodies, respectively. For SARS-CoV, the corresponding numbers were higher and required 200 μg/ml of each antibody to approach 100% inhibition, where 200 μg only achieved 77% and 28% neutralization, respectively, for each individual antibody. These changes in potency and efficacy suggest synergy between CV38-142 and COVA1-16. Synergistic neutralization effects have been analyzed in other viruses (Zwick et al., 2001), including coronaviruses (ter Meulen et al., 2006; Zost et al., 2020), and can be quantified by several algorithms using multiple synergistic models (Ianevski et al., 2017; Wooten and Albert, 2020). Using the most up to date synergy model, our data analysis showed synergistic potency (α>1) between CV38-142 and COVA1-16 in two directions against both SARS-CoV-2 and SARS-CoV pseudoviruses, which suggests reciprocal synergy between CV38-142 and COVA1-16 (Figure S1). Addition of COVA1-16 also improved the maximal efficacy of CV38-142 in neutralizing SARS-CoV as indicated by the positive synergistic efficacy score (β>0) (Figure S1) as well as the neutralization matrix (Figure 2B).

**Figure 2.**
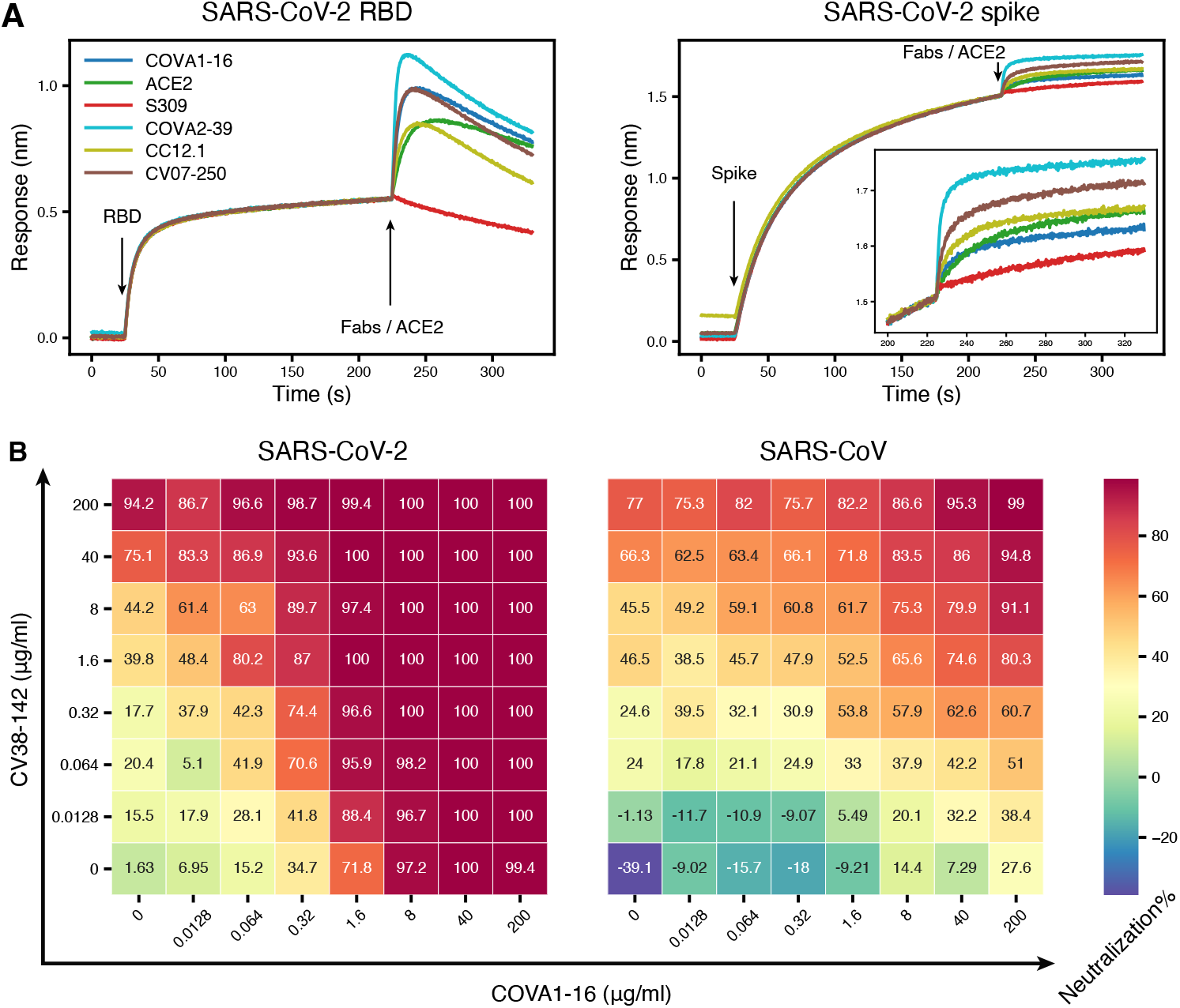
CV38-142 could be combined with antibodies binding to receptor binding site or CR3022 cryptic site. **A.** Competitive binding of CV38-142 to SARS-CoV-2 RBD or spike. Insertion in the right panel shows a zoomed-in view for Fabs/ACE2 binding on spike. A sandwich binding assay was used for the competition assay. CV38-142 IgG was first pre-loaded on the biosensor then SARS-CoV-2 RBD or spike was loaded at the indicated timepoint. The biosensors with captured antibody-antigen complex were tested against binding to a second antibody Fab or human ACE2. Loading events for RBD/spike and the second antibody Fab/ACE2 are indicated by arrows along the timeline (x-axis), while the binding response (nm, y-axis) was recorded in real time as colored lines corresponding to each antibody Fab or ACE2. **B.** Cross-neutralization doseresponse matrix of an antibody cocktail consisting of CV38-142 and COVA1-16. The pseudovirus neutralization assay was performed by addition of mixtures of varying ratios of CV38-142 and COVA1-16. The percentage neutralization for each experiment with SARS-CoV-2 and SARS-CoV is plotted on heatmap matrices with their corresponding color bar shown on the right.

### CV38-142 binds to a proteoglycan site on SARS-CoV-2 RBD

We then determined the crystal structure of SARS-CoV-2 RBD in complex with CV38-142 and COVA1-16 Fabs at 1.94 Å resolution (Figure 3A and Figure S2A, Table S1). COVA1-16 binds to a highly conserved epitope on RBD in the same approach angle as we reported before (Liu et al., 2020). However, CV38-142 binds to a less conserved surface with no overlap with the COVA1-16 epitope (Liu et al., 2020) and involves the N-glycosylation site at N343 on the RBD that is distal to the RBS (Figure 3A and Figure S2A). This N343 glycosylation site is conserved in sarbecoviruses (Figure S3). The crystal structure showed well-resolved density for four of the sugar moieties attached to N343 (Figure S4A). Several hydrogen bonds are made to the glycan from both heavy and light chain (Figure 3B). The V_H_ S100 amide hydrogen bonds to the post-translationally modified N343, and V_H_ R96, V_L_ Y49 and V_L_ S53 hydrogen bond to the core fucose moiety of the glycan as well as water molecules that mediate interactions between CV38-142 and glycan. These interactions contribute to binding between CV38-142 and SARS-CoV-2 RBD as glycan removal from the RBD using PNGase F, or with RBD expressed in HEK293S cells that results in high mannose glycans with no core fucose, results in a decrease in binding to CV38-142 from a K_D_ of 27 nM to 42nM and 168 nM (Figure 3C and Figures S2B–C). Glycan removal resulted in only a slight decrease in binding to SARS-CoV elicited antibody S309 (Figure S2E), which also interacts with the N343 glycan in SARS-CoV-2 RBD (Pinto et al., 2020). To eliminate glycosylation at the N343 site, mutations were introduced into the NxT sequon either at asparagine or threonine residue in both SARS-CoV-2 and SARS-CoV RBDs. An enzyme-linked immunosorbent assay (ELISA) showed a significant drop in binding of CV38-142 to both SARS-CoV-2 and SARS-CoV RBD, while antibody binding to other epitopes, such as CR3022 and CV07-209, were not impacted (Figure S2D). Deep mutational scanning on SARS-CoV-2 RBD previously indicated lower expression of mutants with changes near the glycosylation site, especially at residue 343 (Starr et al., 2020). We therefore used S309 as a probe to show the epitope surface is exposed and can be recognized by S309 (Figure S2D). S309 is less affected by the absence of the N343 glycan as mutation in the NxT sequon at residue 345 had minor impact on S309 binding to the RBD, although there was a significant drop in binding to SARS-CoV-2 RBD N343Q (Figure S2D). Residue 343 also appears to be less tolerant of mutations than residue 345 (Starr et al., 2020). These findings suggest that the complex glycan at N343 (Wang et al., 2020; Watanabe et al., 2020) contributes to RBD binding by CV38-142, especially with its core fucose, rather than simply acting as a glycan shield to antibodies.

**Figure 3.**
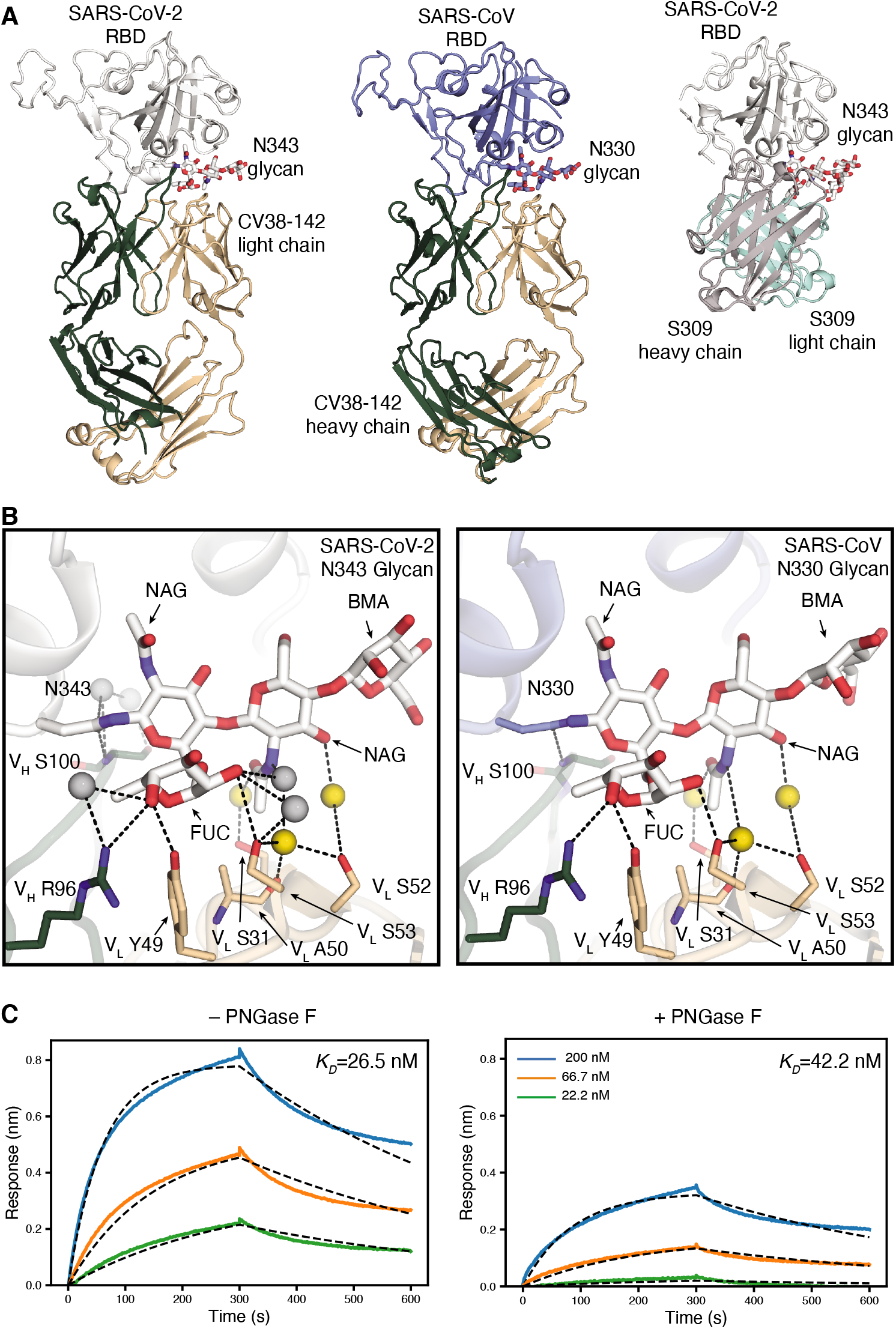
The CV38-142 epitope on the RBD involves an N-glycosylation site on SARS-CoV-2 and SARS-CoV. **A.** Ribbon representation of the crystal structures of SARS-CoV-2 (left) and SARS-CoV (middle) RBD in complex with CV38-142 Fab and comparison to cryo-EM structure of S309 Fab in complex with spike trimer (PDB: 6WPS) (right, only the comparable RBD regions are shown). CV38-142 Fab heavy chain is in forest green and light chain in wheat, S309 Fab heavy chain in grey and light chain in cyan, SARS-CoV-2 RBD in white and SARS-CoV RBD in pale blue. The N343 glycan in SARS-CoV-2 and N330 glycan in SARS-CoV are shown as sticks. The same perspective views are used for the comparison. The overall structure of SARS-CoV-2 RBD in complex with CV38-142 and COVA1-16 is shown in Figure S1A. **B.** Interactions between CV38-142 Fab residues and N343 (SARS-CoV-2) and N330 (SARS-CoV) glycans are shown in stick representation. Water molecules mediating the antibody-antigen interaction are shown in spheres (grey; yellow for shared water-mediated interactions between SARS-CoV-2 and SARS-CoV). Dashed lines (black) represent hydrogen bonds. Residues of the heavy and light chain are both involved in the interactions with glycans. The interactions of CV38-142 with SARS-CoV-2 RBD and SARS-CoV RBD are similar. **C.** Glycan removal in the RBD decreases binding between CV38-142 and SARS-CoV-2 RBD. The binding kinetics were measured by BLI with CV38-142 Fab on the biosensor and RBD in solution. SARS-CoV-2 RBD was pretreated with or without PNGase F digestion in the same concentration and condition before being used in the BLI assay. Concentrations of RBD serial dilution are shown in the right panel. The association and disassociation were recorded in real time (s) in the x axis and response (nm) on the y axis as colored lines. Disassociation constant (K_D_) values were obtained by fitting a 1:1 binding model with fitted curves represented by the dash lines.

In addition to the N343 glycan, interactions with other residues are observed between CV38-142 and SARS-CoV-2 RBD. The V_H_ R58 guanidinium hydrogen bonds to the L441 backbone carbonyl in SARS-CoV-2, while its hydrophobic portion interacts with the alkene region of K444. V_H_ W100c indole hydrogen bonds with the N440 carbonyl and forms a hydrophobic patch with V_H_ V98 and the L441 side chain in SARS-CoV-2 (Figure 4A). The V_H_ S55 backbone carbonyl oxygen hydrogen bonds to the N450 amide (Figure 4A). Besides heavy-chain interactions, the V_L_ Y92 carbonyl oxygen hydrogen bonds to the N440 side chain in SARS-CoV-2 RBD. Overall, CV38-142 interacts with RBD mainly through its heavy chain, which contributes 79% of the buried surface area (BSA) on the RBD (629 Å^2^ out of 791 Å^2^ total BSA as calculated by the PISA program, Figure 4B). Eight polar interactions and two sites of hydrophobic interactions are involved in binding of CV38-142 to SARS-CoV-2 RBD (Table S2).

**Figure 4.**
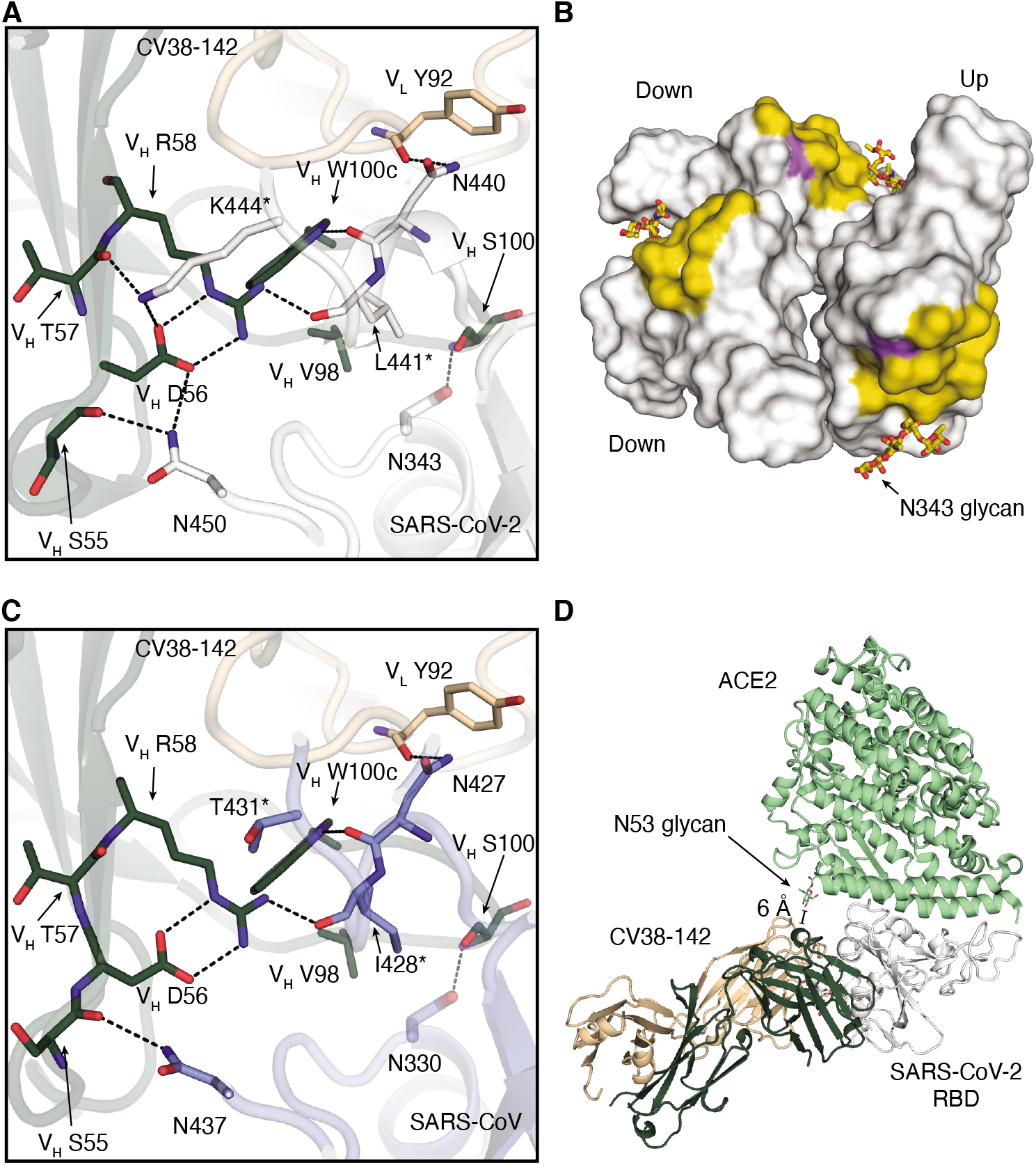
Detailed interactions between CV38-142 and RBDs. SARS-CoV-2 RBD is in white, SARS-CoV RBD in pale blue, CV38-142 heavy chain in forest green and light chain in wheat, and ACE2 in pale green. Corresponding residues that differ between SARS-CoV-2 and SARS-CoV are labelled with asterisks (*). Dashed lines (black) represent hydrogen bonds or salt bridges. **A.** Direct interactions between CV38-142 and SARS-CoV-2 RBD are shown in sticks. **B.** Surface representation of the CV38-142 epitope site in SARS-CoV-2 RBD. The CV38-142 epitope is exposed to solvent regardless of whether the RBD is in the “up” or “down” state. RBDs are shown in surface representation model with symmetry derived from the spike protein (PDB: 6VYB) to show their solvent accessible surface area in either “up” or “down” state. The buried surface area (BSA) was calculated by PISA program (Krissinel and Henrick, 2007). The epitope surface buried by the CV38-142 heavy chain is shown in orange and that by the light chain in purple. The total surface area buried on the RBD by CV38-142 is 792 Å^2^ with 629 Å^2^ (79%) contributed by the heavy chain and 163 Å^2^ (21%) by the light chain. **C.** Direct interactions between CV38-142 and SARS-CoV RBD. The same perspective is used as in **A. D.** Structural alignment illustrating a model with simultaneous binding by CV38-142 and ACE2 to SARS-CoV-2 RBD. Structures of CV38-142 Fab+SARS-CoV-2 RBD and ACE2+SARS-CoV-2 spike are aligned by superimposition of their RBD. The scale bar shows the closest distance between ACE2 and CV38-142, which is 6 A, although some sugars in the N53 glycan are not visible in the electron density map.

### CV38-142 uses a plethora of water-mediated interactions to aid in cross-reactivity with SARS-CoV-2 and SARS-CoV RBDs

The RBD residues involved in CV38-142 interaction with SARS-CoV-2 are not all identical in SARS-CoV RBD (Figure S4). Eight of 20 residues differ in the CV38-142 epitope between SARS-CoV-2 and SARS-CoV. To investigate how CV38-142 accommodates these differences, we determined a crystal structure of CV38-142 Fab in complex with SARS-CoV RBD at 1.53 Å resolution (Figure 3A, Table S1). CV38-142 binds SARS-CoV RBD at the same site with an identical approach angle, albeit interacting with some different residues in the RBD. Interaction with the conserved N330 glycan (Figure 3B and Figure S2D), and the conserved N427 and N437 (Figure 4C) are the same as with SARS-CoV-2. Similar hydrophobic interactions are maintained with I428 in SARS-CoV RBD and L441 in SARS-CoV-2 RBD (Figure 4C). However, interactions with K444 are lost in CV38-142 binding to SARS-CoV RBD due a change to the corresponding T431 in SARS-CoV RBD (Table S2). A hydrophilic surface of CDRH3 of CV38-142 is now juxtaposed to F360 of SARS-CoV RBD compared to S373 of SARS-CoV-2 RBD. The phenyl moiety of F360 adopts heterogeneous conformations with diffuse electron density in the X-ray structure (Figure S4E). Side chains of other epitope residues of SARS-CoV RBD that differ from SARS-CoV-2 RBD are well adapted to the binding interface with no clashes or significant changes in the CV38-142 structure. Thus, the overall binding of CV38-142 to SARS-CoV RBD is essentially identical to SARS-CoV-2 (Figures 1A and 3A) despite a few differences in specific interactions (Figures 4A and 4C). It would appear to be unusual that the binding between an antibody and antigen would be retained at the same level with half of the polar interactions being depleted in the interface of a cross-reacting protein (Table 2). One explanation is the abundance of water molecules mediating interaction between CV38-142 and both SARS-CoV-2 and SARS-CoV RBD. Many conserved water-mediated interactions are found with the peptide backbone in both SARS—CoV-2 and SARS-CoV RBD (Figure 5). The structures here are at high enough resolution to confidently identify these bound water molecules (Figure S4C–D). Water molecules have also been shown to be very important in some other antibody-antigen interfaces (Braden et al., 1995; Wilson and Stanfield, 1993; Yokota et al., 2003). The shape complementarity (SC) (Lawrence and Colman, 1993) between CV38-142 and SARS-CoV-2 or SARS-CoV (0.63 and 0.58, respectively) is lower than for the average for antibody-antigen interactions or protein-protein interactions (Kuroda and Gray, 2016), when water molecules are not considered. Consistent with the SC analysis and high binding affinities, 24 water molecules mediate more than 60 hydrogen bonds between CV38-142 and SARS-CoV-2 RBD (Figure 5A and Figure S4C). A comparable number of water-mediated interactions are also observed with SARS-CoV RBD (Figure 5B and Figure S4D). These water-mediated interactions are mostly conserved in the interaction with CV38-142 with SARS-CoV-2 and SARS-CoV RBDs, with 15 that overlap and mediate interactions with both SARS-CoV-2 and SARS-CoV RBD (Figure 5). Considering the contribution from these water molecules, the loss of some direct contacts between CV38-142 and SARS-CoV RBD may be partially compensated by these abundant water-mediated interactions, suggesting a potential mechanism whereby CV38-142 could resist antigenic drift.

**Figure 5.**
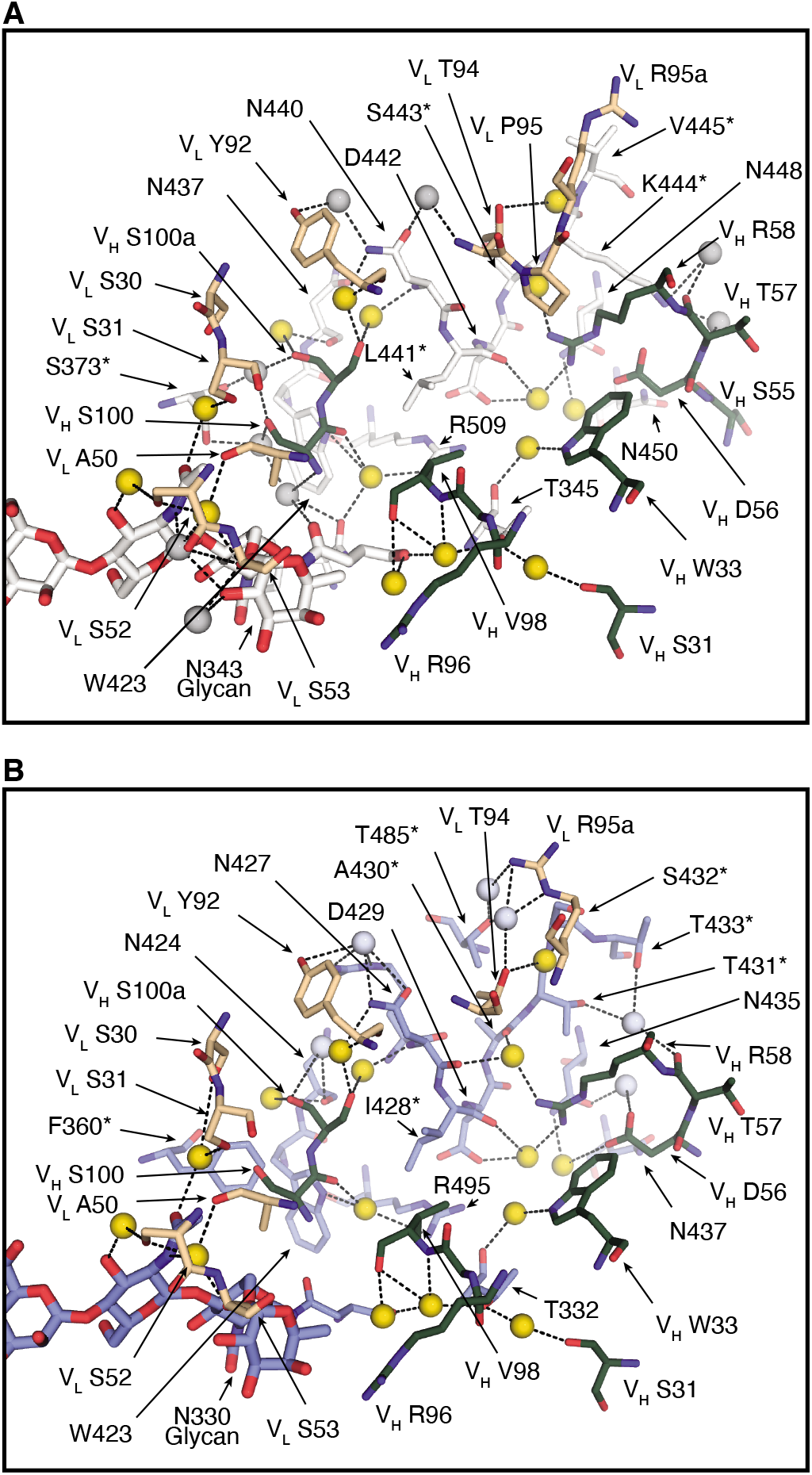
A plethora of water molecules mediating interactions between CV38-142 and SARS-CoV-2 and SARS-CoV RBD. SARS-CoV-2 RBD is in white, SARS-CoV RBD in pale blue, CV38-142 heavy chain in forest green and light chain in wheat. Corresponding residues that differ between SARS-CoV-2 and SARS-CoV are labelled with asterisks (*). Dashed lines (black) represent hydrogen bonds. Amino acid residues as well as the glycans involved in the water-mediated interactions are shown in sticks. Yellow spheres indicate water molecules in the same location in the structures of the CV38-142 Fab+SARS-CoV-2 RBD+COVA1-16 Fab complex (**A**) and the CV38-142 Fab+SARS-CoV RBD (**B**). Grey spheres indicate unique water molecules in each complex structure.

### CV38-142 accommodates rather than competes with ACE2 binding to the RBD

Structure superimposition of ACE2 bound to RBD reveals no clash between ACE2 and CV38-142. The closest distance is 6 Å, which corresponds to the distance between the first NAG moiety of the ACE2 N53 glycan with the H66 imidazole of CV38-142 in the antibody complex with SARS-CoV-2 RBD (Figure 4D). There seems to be sufficient space for the remainder of the glycan to be accommodated due to the large open void between CV38-142 and ACE2. In addition, we observe some flexibility in this region (S60-H66) of CV38-142 that would allow even more room for the ACE2 N53 glycan if both ACE2 and CV38-142 were to bind RBD simultaneously (Figure S5A). BLI sandwich binding assays and the surface plasma resonance (SPR) competition assays revealed that binding of CV38-142 IgG does not occlude ACE2 binding to SARS-CoV-2 RBD or spike protein (Figure 2A and Figure S5B), suggesting no steric block between CV38-142 and ACE2. Since CV38-142 IgG potently neutralize SARS-CoV-2 and SARS-CoV infection in pseudovirus (Figure 1B) and authentic virus assays (Kreye et al., 2020), this finding then poses a question about the mechanism of CV38-142 neutralization of sarbecovirus infection. One explanation is that CV38-142 somehow attenuates ACE2 or other cofactor binding that cannot be observed in the sandwich binding assay or the SPR competition assay. We in fact previously reported that CV38-142 IgG reduced ACE2 binding to SARS-CoV-2 RBD in an enzyme-linked immunosorbent assay (ELISA) by 27% (Kreye et al., 2020). The possible constraint on accommodating the N53 glycan in ACE2 upon simultaneous binding by CV38-142 IgG may contribute to this reduction on ACE2 binding in the ELISA assay (Kreye et al., 2020).

### CV38-142 binds RBD in either “up” or “down” state and could cross-link spikes

Superimposition of the CV38-142 binding epitope onto a cryoEM structure of the spike trimer (PDB: 6VYB) suggests that CV38-142 is capable of binding RBD in both “up” and “down” states (Figure 4B). Consistent with this notion, 2D classification of the negative-stain electron microscopy (nsEM) images reveals that CV38-142 Fab can bind to SARS-CoV-2 or SARS-CoV spikes with various binding stoichiometries (Figure S6A–B). The 3D reconstructions of both SARS-CoV-2 and SARS-CoV spikes indicated that CV38-142 Fab could bind RBDs in either “up” or “down” state (Figure 6A and Figures S6C and S6D). The nsEM reconstructions also showed high flexibility of the RBD that only allowed reconstruction of partial density for the Fab (Figure S6D), suggesting heterogeneous conformations of the RBD when bound with CV38-142 Fab. Since the resolutions of the nsEM data are insufficient to build atomic models of spikes, we fit the crystal structure of CV38-142 Fab+SARS-CoV-2 RBD into the nsEM density map of SARS-CoV-2 spike bound to three CV38-142 Fabs in the two “down”, one “up” state (Figure 6A, pale blue). The tentative fitting model suggests a distance of 88 Å between the heavy chain C-termini of CV38-142 Fabs bound with RBD in “down” state and distances of 146 and 158 Å between the heavy chain C-terminal of CV38-142 Fab bound with RBD in “up” state and one of the RBDs in “down” state (Figure 6B).

**Figure 6.**
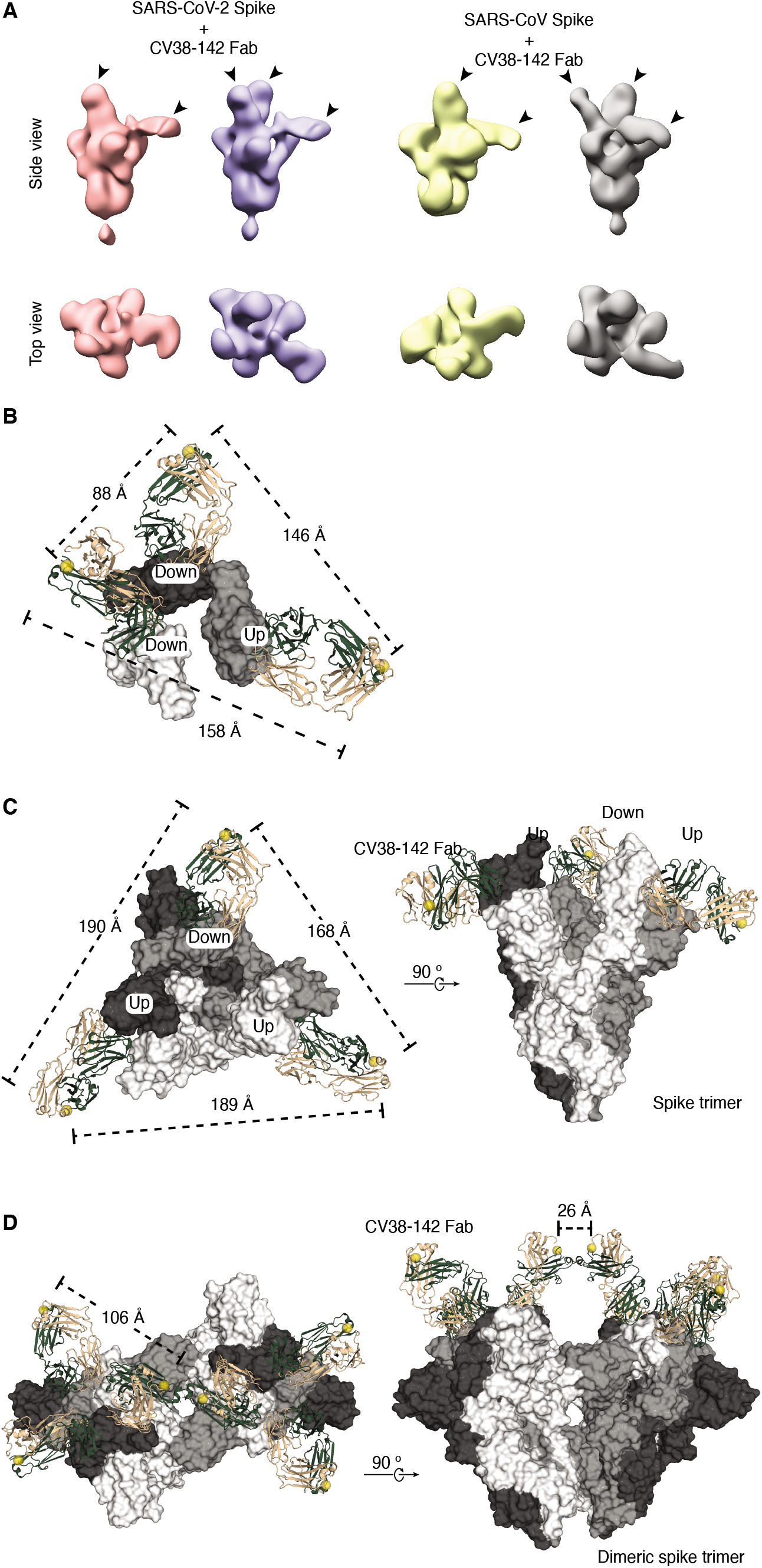
CV38-142 Fab binding to SARS-CoV-2 and SARS-CoV spike trimers. **A.** CV38-142 Fab binding to spike trimers observed by nsEM. Representative 3D nsEM reconstructions are shown of CV38-142 Fab complex with the spike trimers with its RBDs in “up” and “down” states. The location of the bound CV38-142 Fabs are indicated by arrow heads. SARS-CoV-2 (pink) or SARS-CoV (yellow) spikes with at least one “up” RBD and one “down” RBD are bound by two CV38-142 Fabs. The spikes (pale blue to SARS-CoV-2 and grey to SARS-CoV) with RBD in the two “down”, one “up” states are bound by three Fabs. Other binding stoichiometries and conformations are show in Figure S6. **B–D.** C-termini distances of CV38-142 Fab binding to spikes. The three RBDs (**B**) or three protomers (**C–D**) in the spike trimer are shown in white, grey and dark, respectively. CV38-142 Fabs are shown in ribbon representation with heavy chain in forest green and light chain in wheat. The C-termini of CV38-142 heavy chains are shown as spheres (yellow). Dashed lines represent distances among the various combinations of C-termini. **B.** nsEM fitting model. To measure the distances between C-termini of CV38-142 Fabs in nsEM data, the crystal structure of CV38-142 Fab+SARS-CoV-2 was fitted into the nsEM density in A (second from the left). (**C–D**) Structural superimposition of CV38-142 Fabs onto the spike trimer, which is shown in surface representation. Alignment of CV38-142 Fab binding to the spike trimer with RBD in two “up”, one “down” state (PDB: 7CAI) (**C**) or to a dimeric spike trimer that is found in Novavax vaccine candidate NVAX-CoV2373 with RBD in “all-down” state (PDB: 7JJJ) (Bangaru et al., 2020) (**D**). The **B–D** models represent various possibilities of CV38-142 binding to the spike protein on the viral surface.

For spike with RBDs in “2-up-1-down” state, we aligned CV38-142 Fab to the cryoEM structure of SARS-CoV-2 spike (PDB: 7CAI) (Lv et al., 2020). Structural alignment suggests that the C-terminus of the CV38-142 Fab heavy chain points away from the spike center axis due to its particular approach angle (Figure 6C), echoing a similar observation in the nsEM reconstruction data. The distance among the C-termini ranges from 168–190 Å depending on the various combination of RBD states and similar to that measured in the nsEM fitting model (Figure 6B), indicating that it is not possible for an CV38-142 IgG to bind two RBDs bivalently in either two “up” or one “up”, one “down” states within a spike trimer. For spike with RBDs in all “down” state, we aligned the crystal structure of CV38-142 Fab to a cryoEM structure of dimeric spike trimer (PDB: 7JJJ). The structural alignment reveals that the distance between any two C-termini of CV38-142 Fab bound within a spike trimer is around 106 Å, which also suggests that CV38-142 is unlikely to bind two RBDs in the “down” state within a spike trimer (Figure 6D). On the other hand, CV38-142 Fabs can bind RBDs from two adjacent spikes in a dimer seen in Novavax vaccine candidate NVAX-CoV2373 (Bangaru et al., 2020) with a distance of 26 Å between the C-termini of the Fabs, suggesting that a CV38-142 IgG can bind a dimeric spike, or two spikes that are close together, with its two Fabs bound to RBDs from neighboring spikes (Figure 6D). These analyses are in line with the neutralization data, where bivalency plays a critical role on neutralizing both SARS-CoV-2 and SARS-CoV infection as the Fab has much weaker to no inhibition against pseudovirus infection by these sarbecoviruses (Figure 1B).

## DISCUSSION

We report here on a distinct cross-neutralizing epitope in the RBD for an anti-SARS-CoV-2 neutralizing antibody, CV38-142, that cross-reacts with other sarbecoviruses including SARS-CoV-2, SARS-CoV and SARS-related viruses in pangolins and bats (Figure 1A and Figure S3). The epitope of CV38-142 is exposed to solvent regardless of whether RBD in the spike is in either the “up” or “down” states (Figure 4B). A SARS-CoV cross-neutralizing antibody S309, which has been previously characterized, binds to a nearby site and also interacts with the N343 glycan (Pinto et al., 2020). Both CV38-142 and S309 bind to the same face of the RBD to partially overlapping epitopes (Figure 3A and Figure S3) and compete with each other for RBD binding (Figure 2A). However, CV38-142 uses a different approach angle with its heavy and light chain rotated 90° around the epitope and N343 glycan site (N330 in SARS-CoV) compared to S309 (Figure 3A).

Binding of CV38-142 to RBD allows simultaneous binding of RBS antibodies including those encoded by *IGHV3-53* and other germlines (Kreye et al., 2020) as well as others tested in this study. Moreover, we also found that a particular combination of cross-neutralizing antibodies, namely CV38-142 and COVA1-16, to two different sites could synergize to enhance neutralization of both SARS-CoV-2 and SARS-CoV pseudoviruses. The crystal structure of the antibody cocktail in complex with SARS-CoV-2 revealed how two different cross-neutralizing antibodies can interact with the RBD without inhibiting each other (Figure S2). Our neutralization data indicated enhanced potency (i.e. half-maximal inhibitory concentration) and efficacy (maximum percentage of inhibition) with the cross-neutralizing antibody combination (Figure 2B and Figure S1). The improved neutralization may arise from a synergistic effect on trapping the RBD in the up state since binding of COVA1-16 leads the RBD to tilt and twist in the up state (Liu et al., 2020). Since COVA1-16 is representative of cross-neutralizing antibodies that bind to the CR3022 cryptic site (Yuan et al., 2020b), other cross-neutralizing antibodies identified so far (i.e. S304, H014, and EY6A) (Lv et al., 2020; Piccoli et al., 2020; Zhou et al., 2020) that also bind to the CR3022 site (Figure S3) could also be paired with CV38-142 to improve cross-neutralization potency. The receptor binding site is quite diverse in sequence among SARS-CoV-2 and SARS-CoV and already subject to escape mutations; thus, antibodies to cross-neutralizing sites may provide better protection against antigenic drift. Although CV38-142 binds to a less conserved surface of the RBD across sarbecoviruses than COVA1-16, it uses fewer direct contacts and compensates through abundant water-mediated interactions that could accommodate antigenic differences and drift in sarbecoviruses. Given that COVA1-16 has been reported to show resilience to mutations present in the currently circulating variants (Wang et al., 2021a), our study on combinatorial use of cross-neutralizing antibodies provides valuable information to counteract potential escape mutations or antigenic drift in SARS-CoV-2, as well as future zoonotic viruses that could cause threats to global human health.

## ACKNOWLEDGEMENTS

We thank Henry Tien for technical support with the crystallization robot, Jeanne Matteson and Yuanzi Hua for contribution to mammalian cell culture, Wenli Yu for insect cell culture, and Robyn Stanfield for assistance in data collection. We acknowledge BIAFFIN GmbH & Co KG (Kassel, Germany) for the help on the SPR competition assay. We are grateful to the staff of the Advanced Photon Source (APS) Beamline 23ID for assistance. This work was supported by the Bill and Melinda Gates Foundation OPP1170236 and INV-004923 (A.B.W., I.A.W.), NIH R00 AI139445 (N.C.W.) and R01 AI132317 (D.N.), and by the German Research Foundation (H.P.). R.W.S. is a recipient of a Vici fellowship from the Netherlands Organisation for Scientific Research (NWO). This research used resources of the Advanced Photon Source, a U.S. Department of Energy (DOE) Office of Science User Facility, operated for the DOE Office of Science by Argonne National Laboratory under Contract No. DE-AC02-06CH11357. Extraordinary facility operations were supported in part by the DOE Office of Science through the National Virtual Biotechnology Laboratory, a consortium of DOE national laboratories focused on the response to COVID-19, with funding provided by the Coronavirus CARES Act.

## AUTHOR CONTRIBUTIONS

H.L., M.Y., N.C.W., and I.A.W. conceived and designed the study. H.L., M.Y., and C.C.D.L. expressed and purified the proteins for crystallization and binding assay. D.H., L.P. and D.N. provided neutralization data. S.B. and A.B.W. provided nsEM data and reconstructions. H.-C.K., S.M.R., H.P. and J.K. provided CV38-142 antibody sequences and ELISA binding data. M.J.v.G. and R.W.S. provided COVA1-16 antibody sequences. H.L., M.Y. and X.Z. crystallized the antibody-antigen complexes and solved the crystal structures. H.L., M.Y., D.H., S.B., N.C.W., H.-C.K., S.M.R., H.P., J.K., A.B.W. and I.A.W. analyzed the data. H.L., M.Y., N.C.W and I.A.W wrote the paper and all authors reviewed and/or edited the paper.

## DECLARATION OF INTERESTS

A patent application for SARS-CoV-2 antibody CV38-142 was first disclosed in (Kreye et al., 2020) and filed under application number 20182069.3 by some of the authors at Neurodegenerative Diseases (DZNE) and Charité-Universitätsmedizin Berlin. The Amsterdam UMC filed a patent on SARS-CoV-2 antibodies including COVA1-16 under application number 2020-039EP-PR that included the AMC authors on this paper.

## MATERIALS AND METHODS

### Expression and purification of SARS-CoV, SARS-CoV-2 and SARSr-CoV RBDs

The receptor-binding domain (RBD) (residues 319-541) of the SARS-CoV-2 spike (S) protein (GenBank: QHD43416.1), RBD (residues 306-527) of the SARS-CoV S protein (GenBank: ABF65836.1), RBD (residues 315-537) of Guangdong pangolin-CoV (GenBank: QLR06866.1), and RBD (residues 319-541) of Bat-CoV RaTG13 (GenBank: QHR63300.2) were separately cloned into a customized pFastBac vector (Ekiert et al., 2011), and fused with an N-terminal gp67 signal peptide and C-terminal His_6_ tag (Yuan et al., 2020c). Recombinant bacmids encoding each RBDs were generated using the Bac-to-Bac system (Thermo Fisher Scientific) followed by transfection into Sf9 cells using FuGENE HD (Promega) to produce baculoviruses for RBD expression. RBD proteins were expressed in High Five cells (Thermo Fisher Scientific) with suspension culture shaking at 110 r.p.m at 28 °C for 72 hours after the baculovirus transduction at an MOI of 5 to 10. Each supernatant containing RBD proteins were then concentrated using a 10 kDa MW cutoff Centramate cassette (Pall Corporation) followed by affinity chromatography using Ni-NTA resin (QIAGEN) and size exclusion chromatography using a HiLoad Superdex 200 pg column (Cytiva). The purified protein samples were buffer exchanged into 20 mM Tris-HCl pH 7.4 and 150 mM NaCl and concentrated for binding analysis and crystallographic studies.

### Expression and purification of antibodies

Expression plasmids encoding the heavy (HC) and light chains (LC) of the CV38-142 and CV07-250 (Kreye et al., 2020), COVA1-16 and COVA2-39 (Brouwer et al., 2020), CC12.1 (Rogers et al., 2020), and S309 (Pinto et al., 2020) IgG or Fab were transiently co-transfected into ExpiCHO cells at a ratio of 2:1 (HC:LC) using ExpiFectamine™ CHO Reagent (Thermo Fisher Scientific) according to the manufacturer’s instructions. The supernatant was collected at 14 days post-transfection. The IgG antibodies and Fabs were purified with a CaptureSelect^™^ CH1-XL Matrix column (Thermo Fisher Scientific) for affinity purification and a HiLoad Superdex 200 pg column (Cytiva) for size exclusion chromatography. The purified protein samples were buffer exchanged into 20 mM Tris-HCl pH 7.4 and 150 mM NaCl and concentrated for binding analysis, crystallographic studies, negative-stain electron microscopy, and pseudovirus neutralization assays.

### Expression and purification of human ACE2, SARS-CoV-2 RBD and S-HexaPro for binding assay

The N-terminal peptidase domain of human ACE2 (residues 19 to 615, GenBank: BAB40370.1) and the receptor-binding domain (RBD) (residues 319-541) of the SARS-CoV-2 spike (S) protein (GenBank: QHD43416.1) were cloned into phCMV3 vector and fused with C-terminal His-tag. A plasmid encoding stabilized SARS-CoV-2 spike protein S-HexaPro (Hsieh et al., 2020) was a gift from Jason McLellan (Addgene plasmid #154754; http://n2t.net/addgene:154754; RRID: Addgene_154754) and used to express S-HexaPro for the binding assay. The plasmids were transiently transfected into Expi293F cells using ExpiFectamine™ 293 Reagent (Thermo Fisher Scientific) according to the manufacturer’s instructions. The supernatant was collected at 7 days post-transfection. The His-tagged ACE2 or S-HexaPro protein were then purified by affinity purification using Ni Sepharose excel resin (Cytiva) followed by size exclusion chromatography.

### Crystallization and X-ray structure determination

The CV38-142 Fab complexed with SARS-CoV-2 RBD and COVA1-16 Fab (3-mer complex) and CV38-142 Fab complexed with SARS-CoV RBD (2-mer complex) were formed by mixing each of the protein components in an equimolar ratio and incubating overnight at 4°C. 384 conditions of the JCSG Core Suite (Qiagen) were used for setting-up trays for screening the 3-mer complex (12.1 mg/mL) and 2-mer complex (15.0 mg/mL) on our robotic CrystalMation system (Rigaku) at Scripps Research. Crystallization trials were set-up by the vapor diffusion method in sitting drops containing 0.1 μl of protein complex and 0.1 μl of reservoir solution. Crystals appeared on day 3, were harvested on day 7, pre-equilibrated in cryoprotectant containing 15-20% ethylene glycol, and then flash cooled and stored in liquid nitrogen until data collection. Diffraction data were collected at cryogenic temperature (100 K) at beamlines 23-ID-D and 23-ID-B of the Advanced Photon Source (APS) at Argonne National Laboratory and processed with HKL2000 (Otwinowski and Minor, 1997). Diffraction data were collected from crystals grown in drops containing 1.0 M lithium chloride, 10% (w/v) polyethylene glycol 6000, 0.1 M citric acid pH 4.0 for the 3-mer complex and drops containing 0.2 M di-ammonium tartrate, 20% (w/v) polyethylene glycol 3350 for the 2-mer complex. The X-ray structures were solved by molecular replacement (MR) using PHASER (McCoy et al., 2007) with MR models for the RBD and Fab from PDB 7JMW (Liu et al., 2020). Iterative model building and refinement were carried out in COOT (Emsley and Cowtan, 2004) and PHENIX (Adams et al., 2010), respectively. Epitope and paratope residues, as well as their interactions, were identified by using PISA program (Krissinel and Henrick, 2007) with buried surface area (BSA >0 Å^2^) as the criterion.

### Expression and purification of recombinant spike protein for nsEM

The spike constructs used for negative-stain EM contain the mammalian codon-optimized gene encoding residues 1-1208 (SARS-CoV-2, GenBank: QHD43416.1) and 1-1190 (SARS-CoV, GenBank: AFR58672.1) of the spike protein, followed by a C-terminal T4 fibritin trimerization domain, a HRV3C cleavage site, 8x-His tag and a Twin-Strep tags subcloned into the eukaryotic-expression vector pcDNA3.4. For the SARS-CoV-2 spike protein, three amino-acid mutations were introduced into the S1–S2 cleavage site (RRAR to GSAS) to prevent cleavage and two stabilizing proline mutations (K986P and V987P) to the HR1 domain. Residues T883 and V705 were mutated to cysteines to introduce a disulfide for additional S stabilization. For the SARS-CoV spike protein, residues at 968 and 969 were replaced by prolines to generate stable spike proteins as described previously (Kirchdoerfer et al., 2018). The spike plasmids were transfected into 293F cells and supernatant was harvested at 6 days post transfection. Spike proteins were purified by running the supernatant through streptactin columns and then by size exclusion chromatography using Superose 6 increase 10/300 columns (Cytiva). Protein fractions corresponding to the trimeric spike protein were pooled and concentrated.

### nsEM sample preparation and data collection

SARS-CoV-2 and SARS-CoV proteins were complexed with six molar excess of Fab for 1 hour prior to direct deposition onto carbon-coated 400-mesh copper grids. The EM grids were stained with 2 % (w/v) uranyl-formate for 90 seconds immediately following sample application. Grids were either imaged at 120 keV on a Tecnai T12 Spirit using a 4kx4k Eagle CCD. Micrographs were collected using Leginon (Suloway et al., 2005) and the images were transferred to Appion for processing. Particle stacks were generated in Appion (Lander et al., 2009) with particles picked using a difference-of-Gaussians picker (DoG-picker) (Voss et al., 2009). Particle stacks were then transferred to Relion (Zivanov et al., 2018) for 2D classification followed by 3D classification to sort well-behaved classes. Selected 3D classes were auto-refined on Relion and used to illustrate with UCSF Chimera (Pettersen et al., 2004). A published prefusion spike model (PDB: 6Z97) (Huo et al., 2020) was used in our structural analysis.

### Measurement of binding affinities and competition using biolayer interferometry

Binding assays were performed by biolayer interferometry (BLI) using an Octet Red instrument (FortéBio). To determine the binding affinity of CV38-142 Fab with SARS-CoV-2 and SARS-CoV RBDs, 20 μg/mL of His-tagged SARS-CoV or SARS-CoV-2 RBD protein purified from Hi5 cell expression was diluted in kinetics buffer (1x PBS, pH 7.4, 0.002% Tween-20, 0.01% BSA) and loaded on Ni-NTA biosensors (ForteBio) for 300 s. After equilibration in kinetics buffer for 60 s, the biosensors were transferred to wells containing serially diluted Fab samples in running buffer to record the real time association response signal. After a 120 s association step, the biosensors were transferred to wells containing blank running buffer to record the real time disassociation response signal. All steps were performed at 1000 r.p.m. shaking speed. K_D_s were determined using ForteBio Octet CFR software. To determine the binding affinity of CV38-142 Fab or S309 IgG with SARS-CoV-2 RBD pretreated with or without PNGase F, Fab or IgG was loaded on Fab2G or AHC biosensors (ForteBio) for 300 s followed by similar steps to test binding to RBD that was expressed in Expi293F cells. For the sandwich binning assay, CV38-142 IgG was loaded onto AHC biosensors (ForteBio) followed by equilibration in kinetics buffer. The biosensors were transferred to wells containing either SARS-CoV-2 RBD or S-HexaPro proteins in kinetics buffer to allow for antigen association for 200 s followed by testing association of a second antibody Fab or ACE2 for 120 s.

### Measurement of competition using surface plasma resonance

To test whether binding of CV38-142 to SARS-CoV-2 RBD has an impact on the binding of ACE2, a surface plasma resonance (SPR) competition assay was performed on a Biacore T200 instrument (Cytiva) at 25 °C. Biotinylated human ACE2 (residue 18-740, ACROBiosystems) was reversibly immobilized on a CAP sensor chip (Cytiva) using Biotin CAPture Kit (Cytiva). CV38-142 IgG used in the SPR assay was produced in CHO cells and was kindly provided by Miltenyi Biotec, Bergisch Gladbach, Germany. The SPR system was primed and equilibrated with running buffer (10 mM HEPES pH 7.4, 150 mM NaCl, 3 mM EDTA, 0.05% Tween 20) before measurement. 10 nM of SARS-CoV-2 RBD (ACROBiosystems) together with different concentrations of CV38-142 IgG dissolved in the running buffer were injected into the system within 90 s in a flow rate of 30 μl/min followed by a regeneration step between each concentration. The binding response signals were recorded in real time by subtracting from reference cell. And the experiment was repeated once.

### Enzyme-linked immunosorbent assay (ELISA) measuring antibody binding to RBD

Rabbit IgG1 Fc-tagged RBD-SD1 regions of MERS-CoV, SARS-CoV and SARS-CoV-2 as well as point mutants thereof (SARS-CoV: N330Q and T332A, SARS-CoV-2: N343Q and T345A) were expressed in HEK293T cells and immobilized onto 96-well plates as previously described (Kreye et al., 2020). Mutations were introduced by overlap extension PCR and confirmed by Sanger sequencing (LGC Genomics). Human anti-spike RBD monoclonal antibodies were applied at 1 μg/ml and detected using horseradish peroxidase (HRP)-conjugated anti-human IgG (Dianova) and the HRP substrate 1-step Ultra TMB (Thermo Fisher Scientific). HRP-conjugated F(ab’)2 anti-rabbit IgG (Dianova) was used to confirm the presence of immobilized antigens.

### Pseudovirus neutralization assay and synergistic study

Pseudovirus preparation and assay were performed as previously described with minor modifications (Rogers et al., 2020). Pseudovirions were generated by co-transfection of HEK293T cells with plasmids encoding MLV-gag/pol, MLV-CMV-Luciferase, and SARS-CoV-2_Δ18_ spike (GenBank: MN908947) or SARS-CoV spike (GenBank: AFR58672.1). The cell culture supernatant containing SARS-CoV-2 and SARS-CoV S-pseudotyped MLV virions was collected at 48 hours post transfection and stored at −80°C until use. Lentivirus transduced Hela cells expressing hACE2 (GenBank: BAB40370.1) were enriched by fluorescence-activated cell sorting (FACS) using biotinylated SARS-CoV-2 RBD conjugated with streptavidin-Alexa Fluor 647 (Thermo, S32357). Monoclonal antibodies IgG or Fab were serially diluted with DMEM medium supplemented with 10% heat-inactivated FBS, 1% Q-max, and 1% P/S. The serial dilutions were incubated with the pseudotyped viruses at 37°C for 1 hour in 96-well half-well plate (Corning, 3688). After the incubation, 10,000 Hela-hACE2 cells were added to the mixture and supplemented 20 μg/ml Dextran (Sigma, 93556-1G) for enhanced infectivity. The supernatant was removed 48 hours post incubation, and the cells were washed and lysed in luciferase lysis buffer (25 mM Gly-Gly pH 7.8, 15 mM MgSO4, 4 mM EGTA, 1% V/V Triton X-100). After addition of Bright-Glo (Promega, PR-E2620) according to the manufacturer’s instruction, luminescence signal was measured in duplicate. At least two biological replicates were performed for each assay. The IgG half-maximal inhibitory concentration (IC_50_) values were calculated using “One Site - Fit LogIC50” regression in GraphPad Prism 9. For synergy assessment of two monoclonal antibodies, an antibody cocktail matrix was prepared by a combination of mixing a fixed concentration of CV38-142 and increasing the concentration of COVA1-16 or increasing the concentration of CV38-142 and fixing the concentration of COVA1-16. Neutralization percentages for each combination were measured and calculated the same way as the pseudovirus neutralization assay. The neutralization data were converted to the input format for the synergy program (Wooten and Albert, 2020). Synergy scores were calculated by fitting the multidimensional synergy of combinations (MuSyC) model, which is a synergy model based on a multidimensional extension of the Hill equation that allows non-linear dose-response surface contour (Meyer et al., 2019). MuSyC model quantifies synergy in bidirectional way and distinguishes synergies between potency and efficacy. The synergy parameter α_12_, namely synergistic potency quantifies how the second antibody changes the first’s potency and α_21_ quantifies how the first changes the second’s potency. The MuSyC model fitting with the synergy program also gives two other parameters, namely synergistic efficacy (β) and synergistic cooperativity (γ) score (Wooten and Albert, 2020). The β score denotes synergistic efficacy, which quantifies the percent change on the maximal efficacy of the antibody combination compared to the most efficacious single agent. The γ_12_ score denotes how the first antibody changes the second’s Hill slop while γ_21_ denotes how the second changes the first’s Hill slop.

### Shape complementarity analysis

Shape complementarity values (Sc) were calculated using SC program as described previously

(Lawrence and Colman, 1993).

---

**Figure S1.**
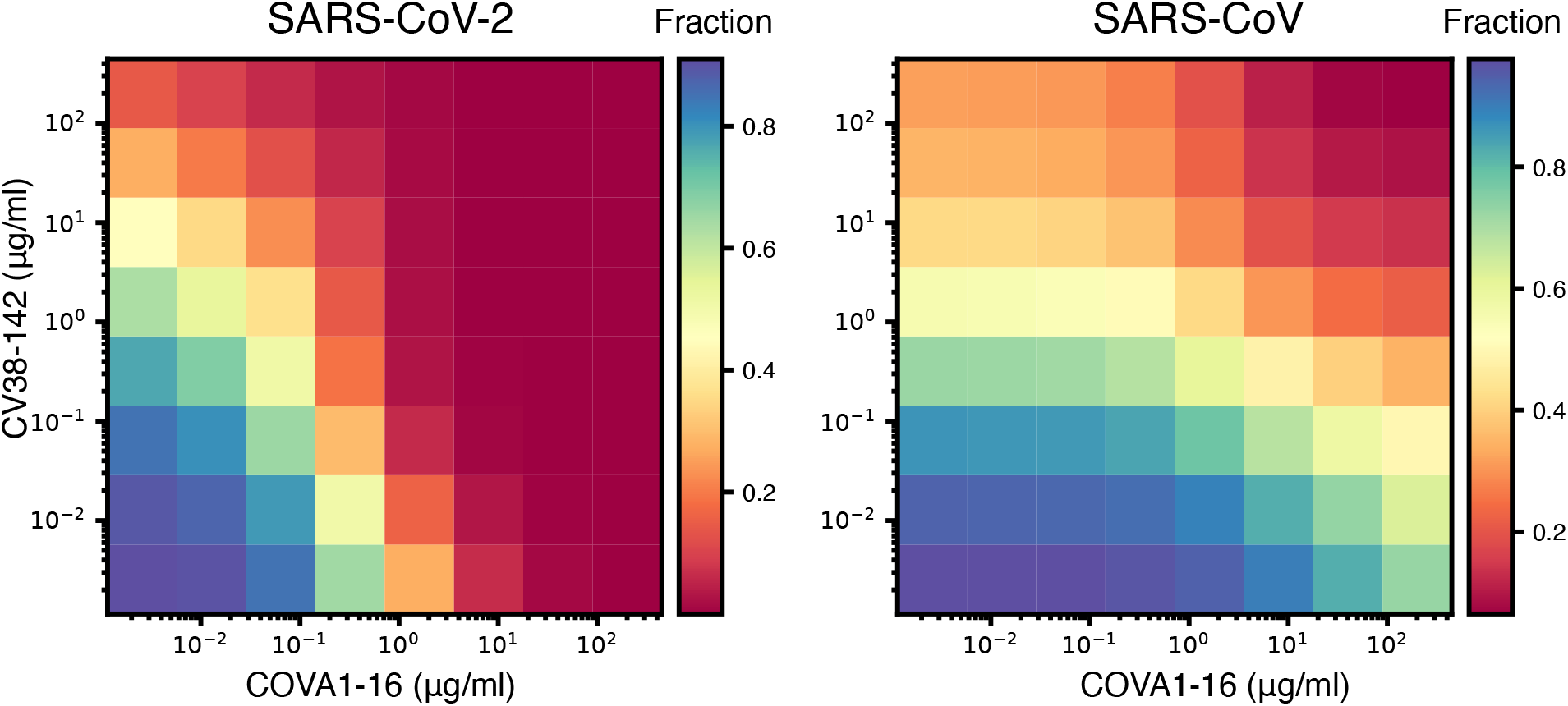
Quantification of synergy between CV38-142 and COVA1-16 using the MuSyC model. Neutralization percentage was used to generate the fraction data with 1 indicates no neutralization and 0 indicates 100% neutralization. Heatmap plot shows the fraction data used for synergy quantification. A>1, γ>1, or β>0 indicate synergism while α<1, γ<1, or β<0 indicate antagonism. CV38-142 was assigned as the first antibody and COVA1-16 was assigned as the second antibody in the analysis. For SARS-CoV-2, CV38-142 and COVA1-16 synergistically change each other’s neutralization potency (α_21_=5314, α_12_=671) and CV38-142 increase the steepness of COVA1-16’s neutralization Hill slope, while COVA1-16 decrease the steepness of CV38-142’s neutralization Hill slop (γ_21_= 2.1, γ_12_=0.38). For SARS-CoV, CV38-142 and COVA1-16 synergistically change each other’s neutralization potency (α_21_=27, α_12_=123) and COVA1-16 increased the efficacy of CV38-142 as indicated by the positive synergistic efficacy score (β=0.4). However, the synergistic efficacy (β) in SARS-CoV-2 neutralization and synergistic cooperativity (γ) in SARS-CoV neutralization are ambiguous (not interpretable) at a 95% confidence interval.

**Figure S2.**
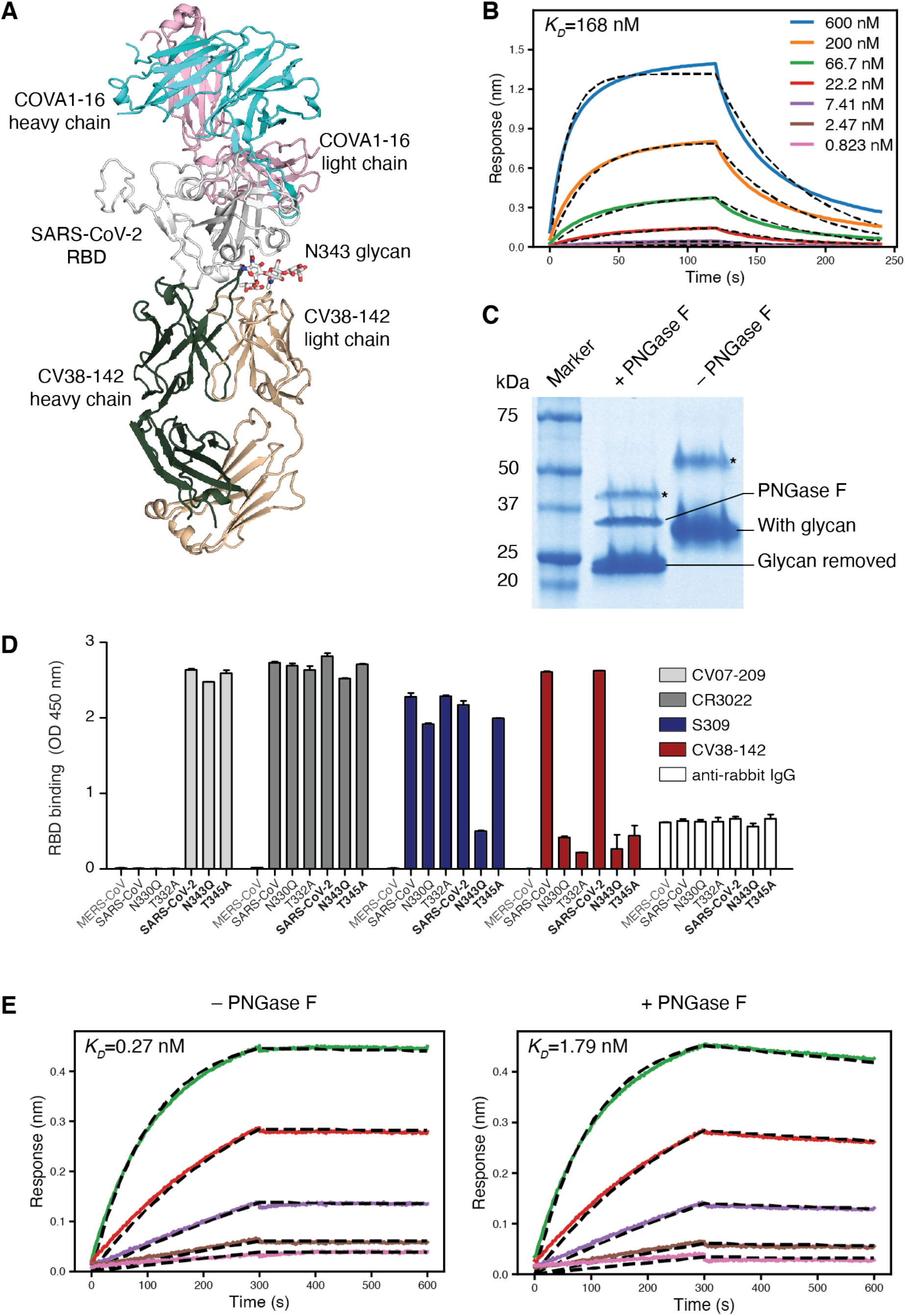
N343 glycan involved in binding to CV38-142. A. Crystal structures of SARS-CoV-2 RBD in complex with CV38-142 and COVA1-16 Fabs. Ribbon representation of SARS-CoV-2 complexed with both CV38-142 Fab and COVA1-16 Fab. The N343 glycan is shown in sticks. SARS-CoV-2 RBD is in white, CV38-142 heavy chain in forest green and light chain in wheat, and COVA1-16 heavy chain in cyan and light chain in pink. There is no overlap between COVA1-16 and CV38-142 epitope as well as no interaction between COVA1-16 Fab and CV38-142 Fab when bound to the same RBD. **B.** Decreased binding affinity between CV38-142 Fab and SARS-CoV-2 RBD expressed in HEK293S cell. HEK293S cell does not have N-acetylglucosaminyltransferase I (GnTI) and therefore protein expressed in this cell lack complex N-glycans (Reeves et al., 2002). N343 glycan of SARS-CoV-2 RBD expressed in HEK293S cell has no fucose moiety and abolishes its interaction to CV38-142 as shown in Figure 3B. Binding kinetics were measured by biolayer interferometry (BLI) with RBDs on the biosensor and Fab in solution. Concentrations of Fab serial dilution are shown in upper right insert. The association and disassociation were recorded in real time (s) on the x axis with binding response (nm) on the y axis with colored lines. Disassociation constant (K_D_) values were obtained by fitting a 1:1 binding model. The fitted curves are represented by the dash lines (black). **C.** PNGase F treatment removes glycans in the SARS-CoV-2 RBD. Non-reducing sodium dodecyl sulphate-polyacrylamide gel electrophoresis (SDS-PAGE) showed the shifted bands between treated and untreated SARS-CoV-2 RBD. Lanes of protein marker, PNGase F treated sample, control sample are indicated above the gel. Protein bands corresponding to SARS-CoV-2 RBD with glycan, with glycan removal, and PNGase F are labeled. Asterisk (*) indicates a small fraction of dimeric RBD formed during protein production. **D.** Mutation in the N343 sequon results in a large decrease in CV38-142 binding to the SARS-CoV-2 RBD. Rabbit IgG1 Fc-tagged RBDs of MERS-CoV, SARS-CoV, SARS-CoV-2 as well as mutant RBDs were coated on a 96-well plate. Binding of the indicated anti-RBD antibodies was tested using an enzyme-linked immunosorbent assay (ELISA). Abolishing the N343 glycosylation by introducing either N343Q or T345A in SARS-CoV-2 and N330Q or T332A in SARS-CoV RBD significantly decreased the CV38-142 binding, with no obvious loss on binding by other antibodies such as CR3022 and CV07-209. S309 appears to be less susceptible to the absence of N343 glycosylation. Note that S309 binds T345A stronger than N343Q, although both mutations lead to no glycosylation at residue 343. N343Q in SARS-CoV-2 may either lead to some steric clashes for antibody binding to the N343Q site that is not the case for T345A or result in a less stable RBD that interferes with the binding detection, as seen by deep mutational scanning (Starr et al., 2020). Two independent repeats were performed, and bar values indicate mean RBD binding with error bars represent the standard deviation. **E.** N343 glycan aids S309 binding to SARS-CoV-2 RBD (Pinto et al., 2020). The binding kinetics were measured by BLI with S309 on the biosensor and RBD in solution. SARS-CoV-2 RBD was treated with or without PNGase F digestion in the same concentration and condition before being used in the BLI assay. Concentrations of RBD serial dilution are color coded as in **B**. The association and disassociation were recorded in real time (s) on the x axis and response (nm) on the y axis as colored lines. Disassociation constant (K_D_) values were obtained by fitting a 1:1 binding model with fitted curves represented by the dash lines.

**Figure S3.**
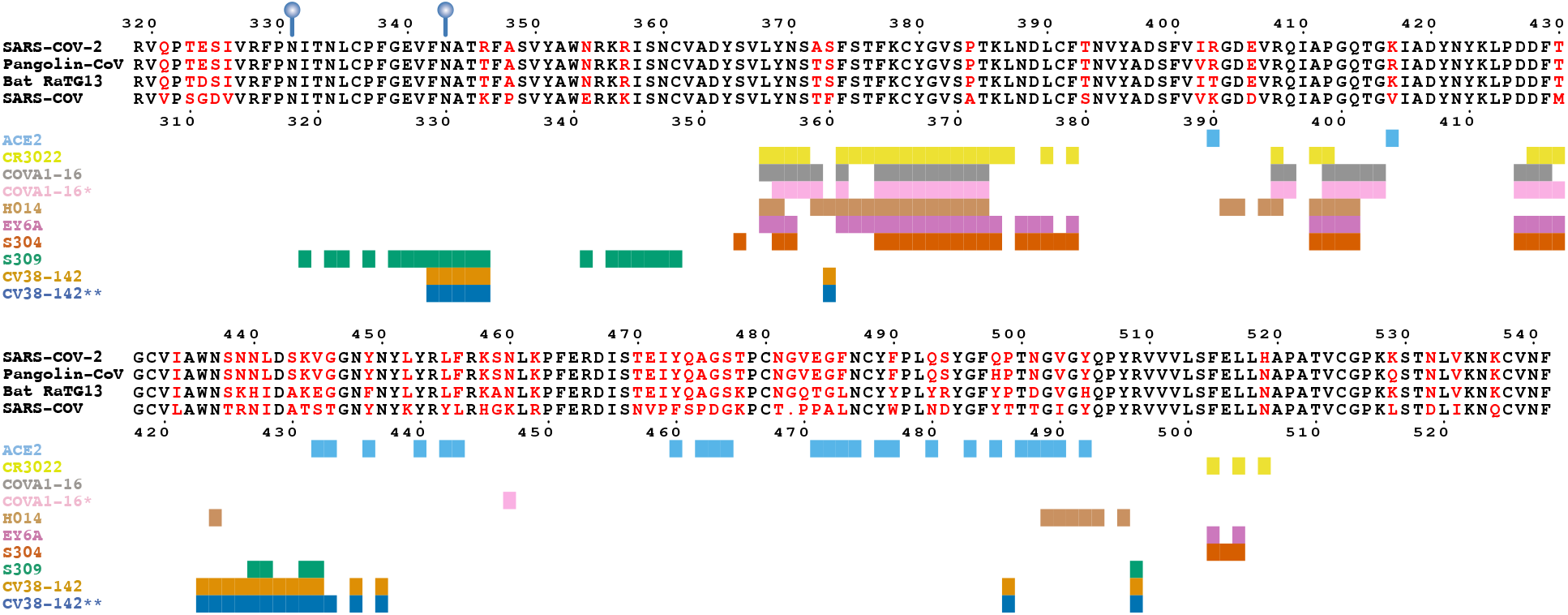
CV38-142 epitope and comparison with other cross-reactive antibodies. Epitopes of cross-reactive antibodies on SARS-CoV-2 or SARS-CoV RBD. Sequence alignment of CV38-142 reactive RBDs from SARS-CoV-2, SARS-CoV, bat coronavirus RaTG13, and Guangdong pangolin coronavirus RBD with non-conserved residues highlighted in red. The conserved glycosylation sites are marked with blue balloons. Numbers corresponding to SARS-CoV-2 RBD and SARS-CoV RBD are labelled every ten residues above and below the sequences panel. Colored bars representing the RBD epitope residues corresponding to each antibody or ACE2 are shown under the sequence panel with their ligand name (ACE2 or antibody) on the left. Epitope residues or ACE2-interacting residues are assigned as BSA>0 Å^2^ as calculated by the PISA program (Krissinel and Henrick, 2007) for SARS-CoV-2 RBD with ACE2 (PDB: 6M0J), CR3022(PDB: 6XC3), COVA1-16(PDB: 7JMW), H014(PDB: 7CAH), EY6A(PDB: 6ZCZ), S304 and S309 (PDB: 7JX3). * indicates COVA1-16 epitope on SARS-CoV-2 RBD calculated from its structure complexed with CV38-142 Fab and SARS-CoV-2 RBD reported in this study as compared to that without CV38-142 (above). The slight discrepancy in COVA1-16 epitope residues is due to the improvement in resolution rather than the simultaneous binding of CV38-142. ** indicates CV-38-142 epitope on SARS-CoV RBD reported in this study in comparison to that on SARS-CoV-2 RBD also reported in this study (above).

**Figure S4.**
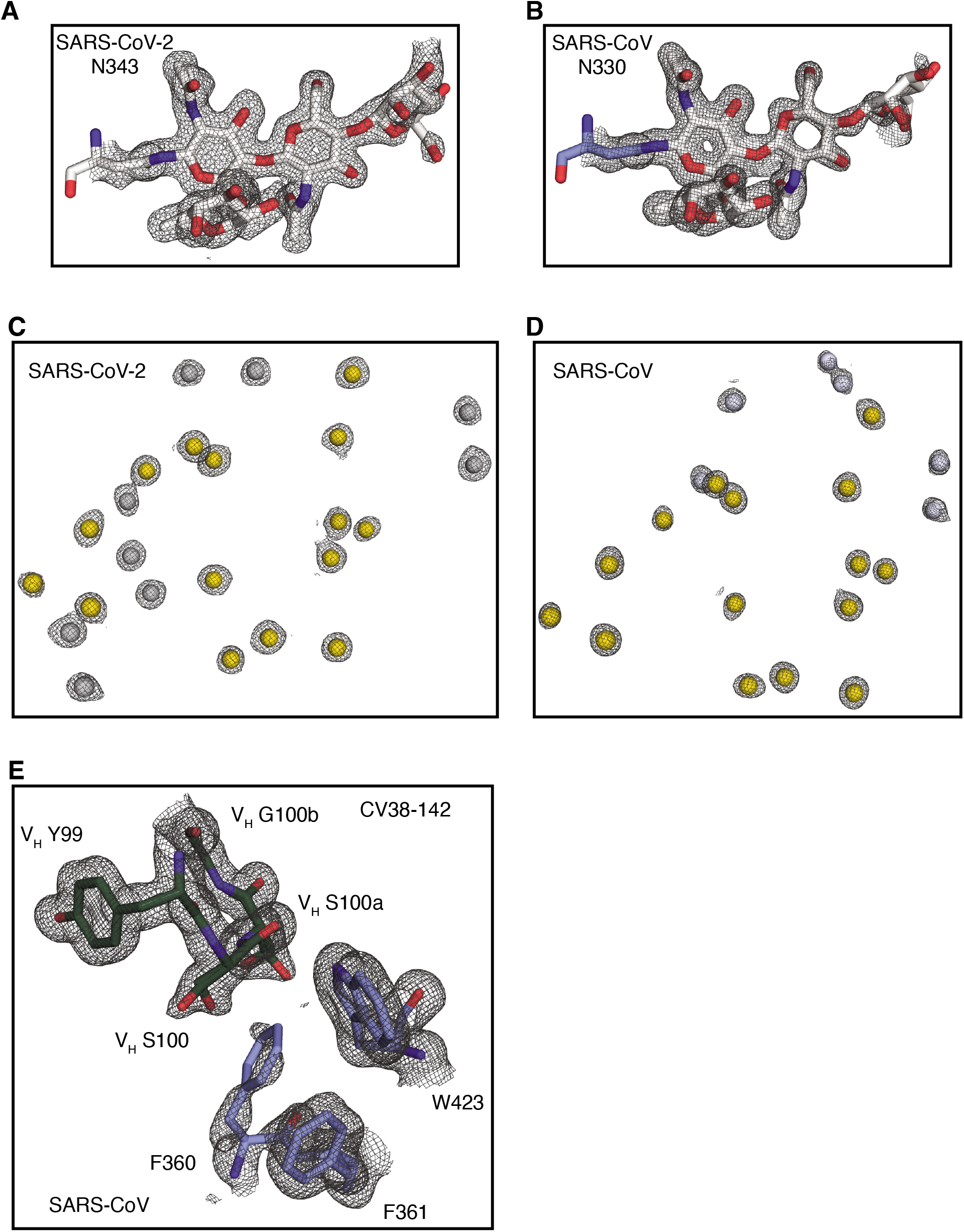
Electron density for glycans, water molecules and region around F360 in the RBD. 2mFo-DFc Sigma-A weighted maps were calculated by Phenix software and contoured at 1.0 σ to show electron density in mesh with the refined structure in spheres (water molecules) or sticks. Glycans and residues are shown in sticks. Water molecules are shown in spheres. Maps are shown in grey meshes. **A.** Electron density for the SARS-CoV-2 N343 glycan. **B.** Electron density for the SARS-CoV N330 glycan. **C.** Electron density for waters in the interface between CV38-142 and SARS-CoV-2 RBD. Shared waters interacting to both SARS-CoV-2 and SARS-CoV RBD are highlighted in yellow. **D.** Electron density for waters in the interface between CV38142 and SARS-CoV RBD. **E.** Electron density for F360 of SARS-CoV RBD and its surrounding residues. CV38-142 is in forest green and SARS-CoV RBD in pale blue.

**Figure S5.**
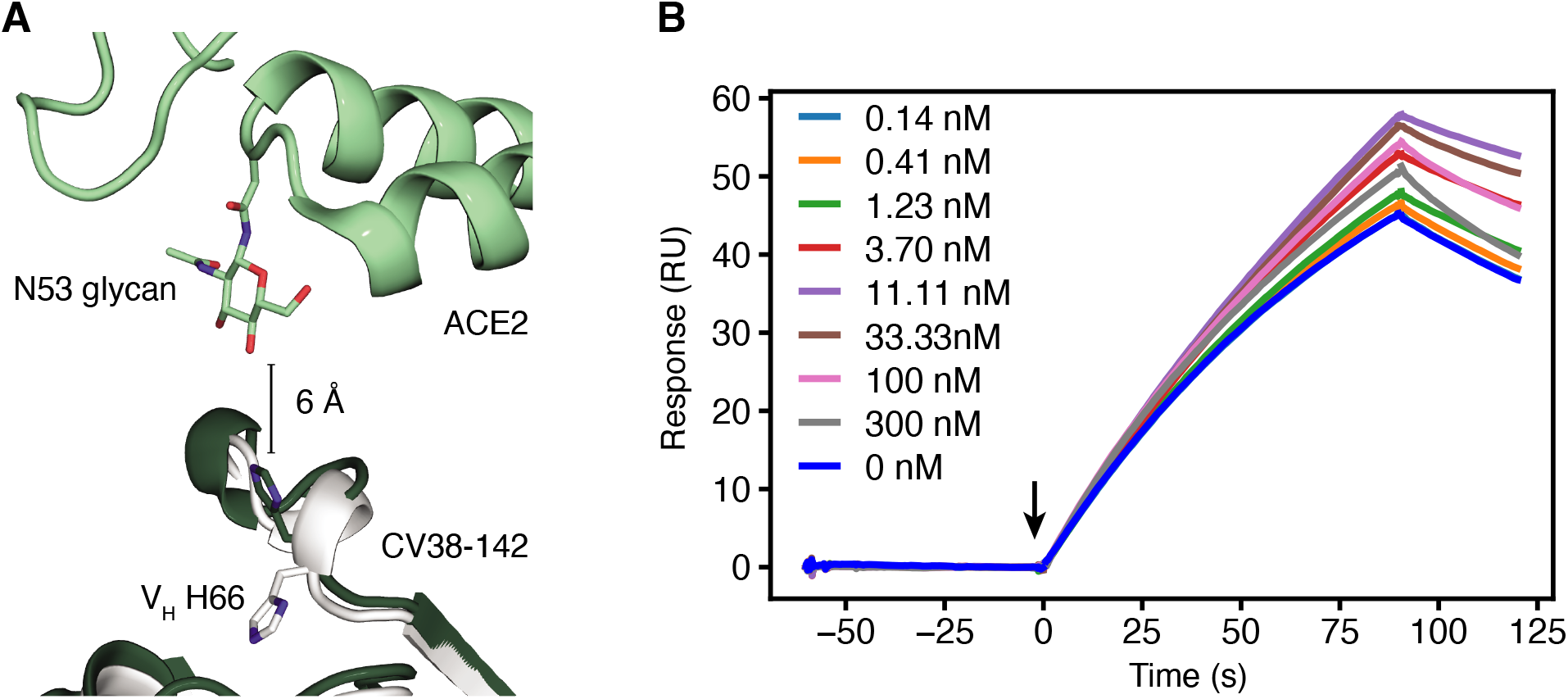
**A.** Close-up view of S60-H66 region of the CV38-142 Fab that is close to the N53 glycan of ACE2 if both were to bind SARS-CoV-2 RBD. Structures of CV38-142 Fab+SARS-CoV-2 RBD (forest green), CV38-142 Fab +SARS-CoV RBD (white), and ACE2+SARS-CoV-2 spike are aligned by superimposition of their RBD. V_H_ H66 of CV38-142 in the Fab complex with SARS-CoV-2 and SARS-CoV RBD is shown as sticks. The closest distance between ACE2 and CV38142 is 6 A, while V_H_ H66 as well as the rest of region S60-H66 of CV38-142 show some flexibility to accommodate the N53 glycan of ACE2. **B.** Surface plasma resonance (SPR) competition assay. Human ACE2 was immobilized on a CAP sensor chip before the measurement of competition. Binding to ACE2 was monitored in real time. Arrow indicates the timepoint of injection of SARS-CoV-2 RBD+CV38-142 IgG mixture. The concentration of SARS-CoV-2 RBD is fixed to 10 nM with while increasing the concentration of CV38-142 IgG as indicated in the insert legend. No inhibition from CV38-142 IgG has been observed as all the concentration tested give very similar on- and off-rate in the binding of ACE2 to the given concentration of SARS-CoV-2 RBD.

**Figure S6.**
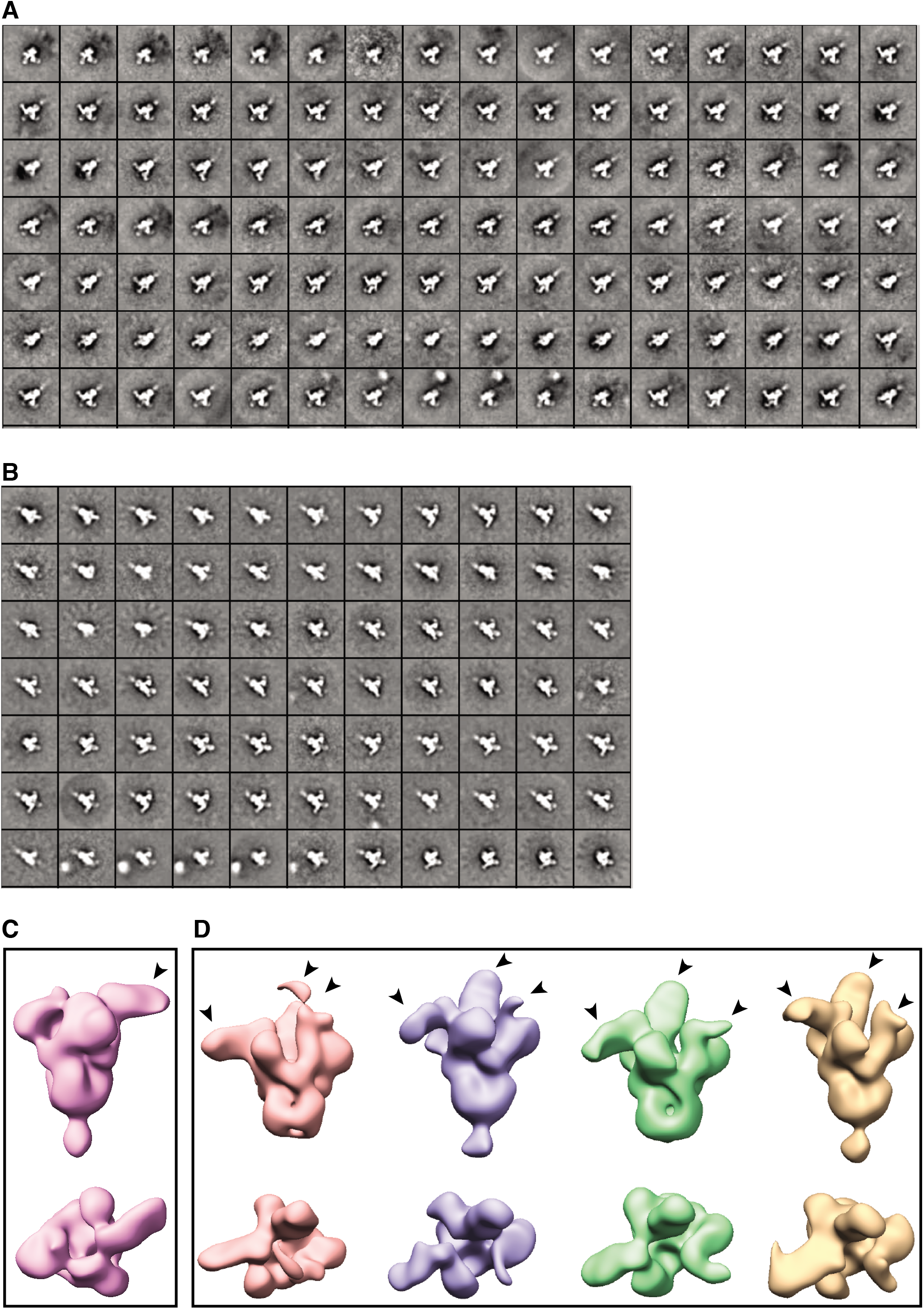
nsEM 2D images and 3D reconstruction of CV38-142 Fab in complex with SARS-CoV-2 and SARS-CoV spikes. **A-B.** 2D classification of the nsEM images showing various binding stoichiometries between CV38-142 Fab and SARS-CoV-2 spike (**A**) and SARS-CoV spike (**B**). **C-D.** 3D reconstruction of SARS-CoV spike bound with one CV38-142 Fab (**C**) and three CV38-142 Fabs (**D**). Arrow heads indicate the RBD with Fab bound. **C.** The spike with at least one “up” RBD. CV38-142 Fab binds to the RBD in the “up” conformation. **D.** The spike with at least one “up” RBD and one “down” RBD. Fabs show binding at various angles among these 3D reconstructions due to flexibility of the RBD in the spike and whether the RBD is up or down. The EM maps for some RBDs bound to Fab are difficult to interpret due to the heterogeneous conformations resulting from the RBD flexibility.

**Table S1.**
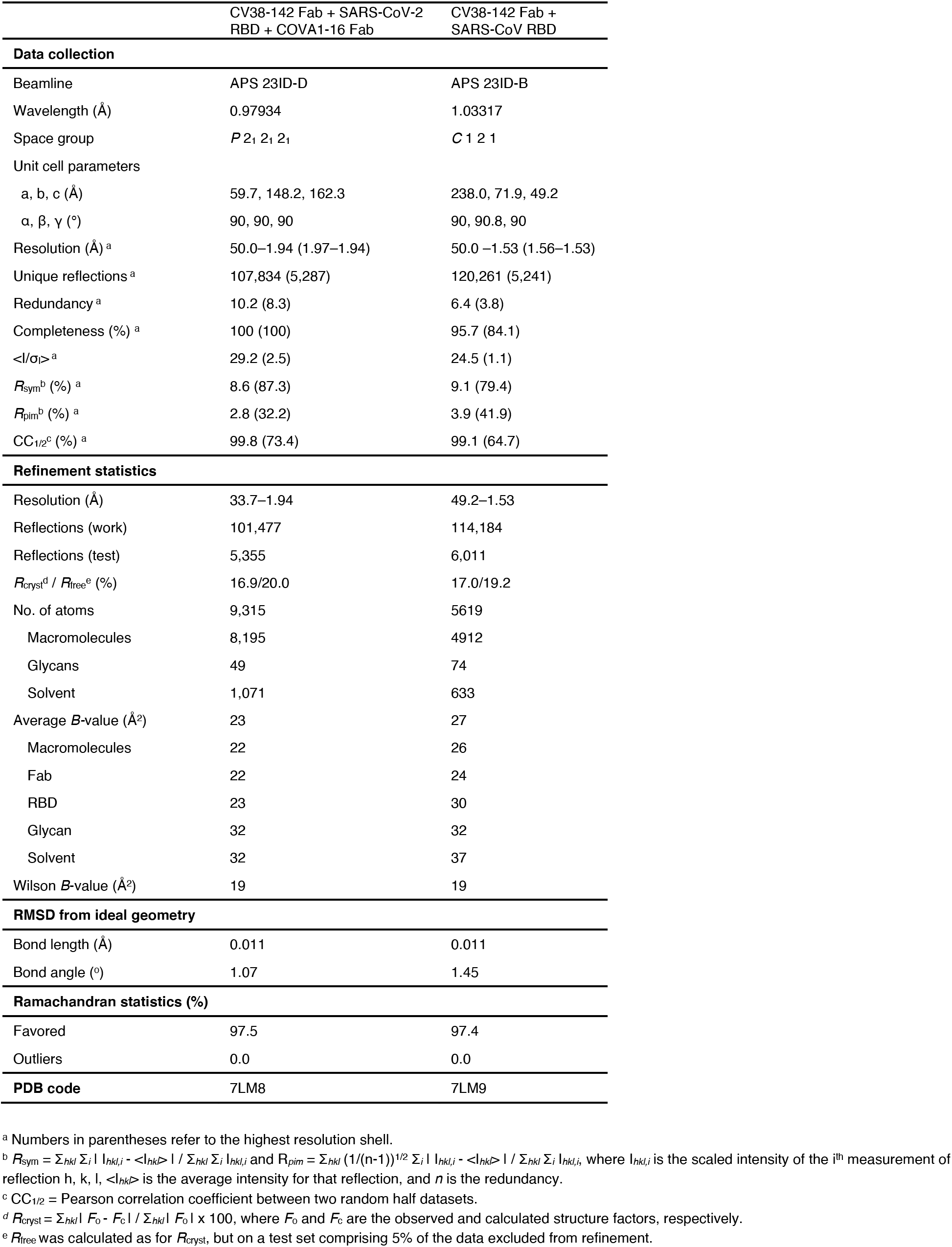
Crystallographic data collection and refinement statistics.

**Table S2.**
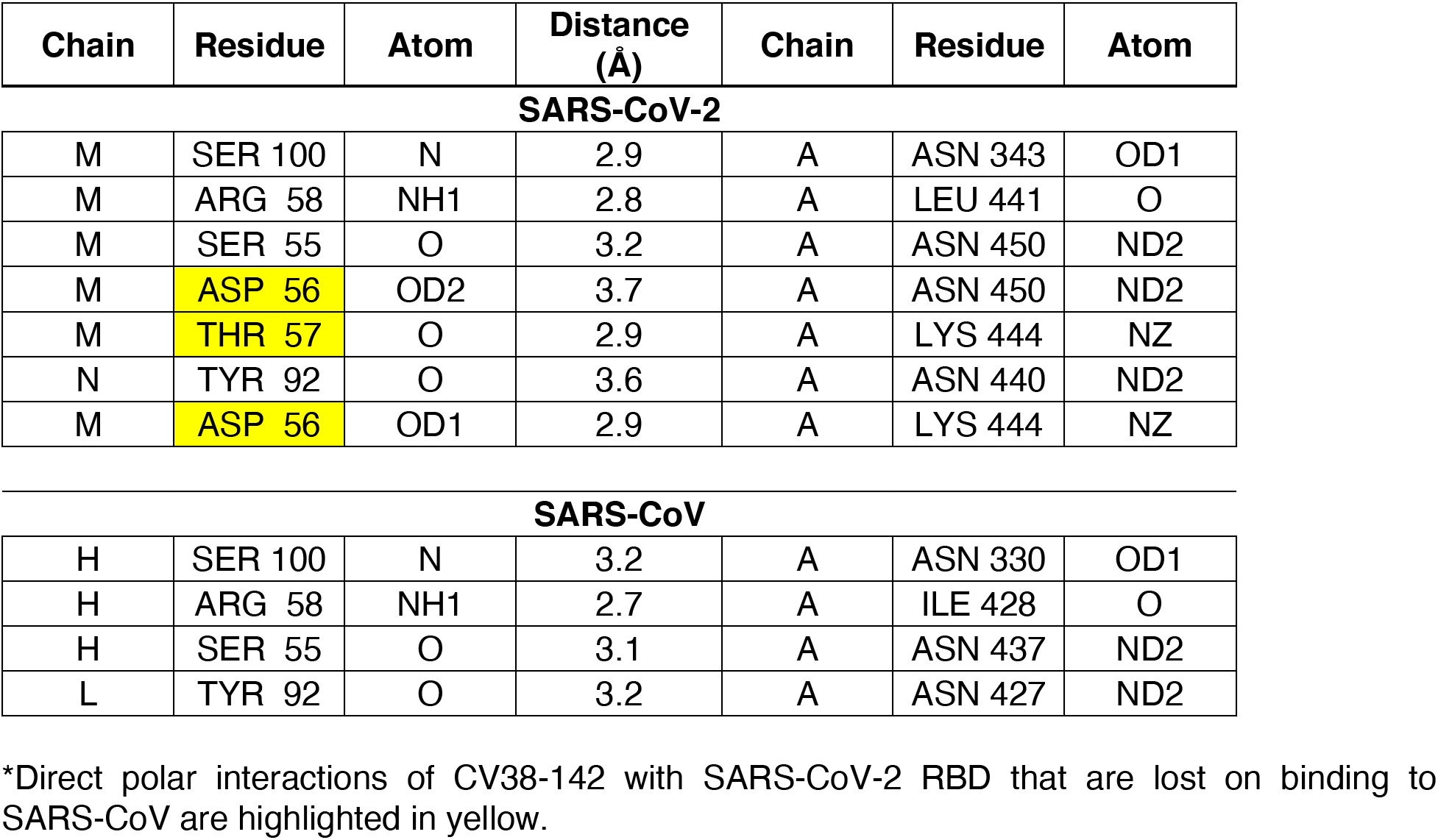
Polar interactions identified at the antibody-antigen interface using the PISA program*.

